# Potential impact of climate change on Nearctic leafhopper distribution and richness in North America

**DOI:** 10.1101/2023.12.13.571535

**Authors:** Abraão Almeida Santos, Jordanne Jacques, Edel Pérez-López

## Abstract

Climate change significantly contributes to shifts in the geographical range of pests and diseases. Leafhoppers (Hemiptera: Cicadellidae), known vectors of phytoplasmas pathogens, are linked to the transmission of more than 600 diseases affecting a thousand plant species worldwide. Despite this, the potential effects of climate change on leafhopper vectors of phytoplasmas remain a critical knowledge gap. To address this gap, our study investigated the potential impact of climate change on 14 species of Nearctic leafhoppers previously associated with phytoplasma-related diseases. Using the MaxEnt species distribution algorithm and other ecological niche modeling techniques, we assessed *(i)* the expected species richness under current climate conditions and four future scenarios and *(ii)* the environmental niche similarity among these species across these scenarios. Our projections suggest that the eastern region of North America holds the potential for the highest species richness, a trend expected to persist across all future scenarios, gradually expanding eastward. Notably, our findings indicate the increasing suitability of northern Canada for more species. Network analysis further revealed a remarkable similarity in environmental niches among most leafhopper species. Moreover, across the four future scenarios, there is a tendency for an increase in this similarity. Altogether, our study underscores the potential persistent presence of Nearctic leafhoppers in their current habitats while pointing to a shift toward northern North America in future scenarios. These findings have significant implications for sustainable pest management practices, prompting a necessary discussion on strategies to mitigate climate change and pest migration’s impact on agricultural systems.

## INTRODUCTION

Climate change significantly contributes to alterations in species distributions, prompting shifts in their geographic ranges^1,2^. Over the past six decades, crop pests and diseases have notably moved toward northern latitudes globally, progressing at approximately 2.7 km annually since 1960^3^. This phenomenon is primarily attributed to global warming and trade dynamics^4,5^. However, the response of these organisms to the ongoing environmental changes is intricate and conditional upon various determinants, notably their geographic origins^6–8^. Tropical pest species, for instance, typically exhibit a restricted tolerance to temperature variation and tend to live close to their maximum thermal limits^6,8^. In contrast, temperate species contend with a broader spectrum of climate extremities, including temperature fluctuations^6^. As a result, projections suggest a potential reduction in the suitable habitats for tropical pest species with the anticipated progression of climate warming^6,9^. Conversely, a favorable trend is expected for temperate species, given their adaptations to adapt to more diverse climatic events^4,6^.

Recently, the efforts to assess the impacts of climate change on biological systems have intensified^10^. Species distribution modeling (SDM), also known as ecological niche modeling (ENM), has risen as a crucial tool to explore the potential effects of climate change on the geographic distribution of organisms^11^. However, within entomology, research efforts have predominantly centered around specific orders, such as Lepidoptera and Diptera, while less attention has been dedicated to others like Hemiptera^10^. This bias may stem from the notorious role Lepidoptera plays in global research and the importance of Diptera as disease vectors for vertebrates^10^. However, Hemiptera also includes significant plant disease vectors, causing substantial damage worldwide^12^. To gain a broader understanding of climate change’s impact on insect distribution, particularly those acting as plant disease vectors, increased research focus on understudied insect orders like Hemiptera is essential.

Leafhoppers (Hemiptera: Cicadellidae) constitute a herbivorous group known for their varied feeding habits across vascular plants^13,14^. With an estimated 21,351 species described globally^15^, these insects exhibit a remarkable characteristic: about 90% of their genera are endemic to specific geographical regions^16^. The Nearctic region, spanning from Mexico to the Arctic, stands out as a leafhopper hotspot, with nearly 3,000 described species showcasing a high level of endemism and diversity^14–20^. Nearctic leafhoppers are well-adapted to various climatic conditions, surviving from harsh winters to scorching summers, finely synchronizing their life cycles with these climatic patterns^14,17–19^. This adaptation suggests that environmental factors such as temperature and continental winds could primarily influence their distribution.

Increasing temperatures favor Nearctic population growth, as higher temperatures reduce the species’ life cycle duration^21,22^. Additionally, continental winds facilitate their dispersal into northern regions during the spring and summer^23–25^. Although the distribution of many leafhopper species is unknown or understudied in several areas of the Nearctic region, it could be plausible to anticipate that ongoing global warming and other climatic changes may offer Nearctic leafhoppers opportunities to expand their potential geographical range.

While leafhopper herbivory can damage leaves, causing necrosis, drops, stunting, smaller seeds, and reduced photosynthesis, leading to decreased yield (e.g., ^26^), their main recognized role is transmitting plant pathogenic viruses and bacteria^12,27^. Phytoplasmas, for instance, are pathogens primarily disseminated by leafhoppers^12^. This group of phloem-limited obligatory parasites belongs to the provisional genus ‘*Candidatus* Phytoplasma’ within *Mollicutes* class^27^. Over 100 leafhopper species carry phytoplasmas linked to more than 600 diseases affecting a thousand plant species worldwide^12,27–29^. Typical symptoms of phytoplasmas diseases include phyllody, virescence, stunt, witches’ broom, fruits appearing as green flowers, and small, curly, and red leaves^22,30^. Examples of such economically damaging diseases in commercial crops include aster yellows in canola^31^, bushy stunt in corn^32^, false blossom in berries^33,34^, green petal in strawberries^22^, and grapevine yellows (*Flavescence dorée* and *Bois noir*) in grapevines^35,36^.

In the Nearctic region, leafhoppers exhibit oligophagous or polyphagous feeding behaviors, demonstrating a broad consumption pattern across diverse host plants that can serve as main hosts or reservoirs of phytoplasmas^14,17,19,37^. In this region, 38 leafhopper species have been identified as vectors or suspected vectors of phytoplasmas affecting horticultural, fruit, and field crops^33,34,38–40^. However, recent findings by our group indicate a tripling incidence of phytoplasma diseases in the Nearctic over the past decade^30^. This trend suggests several potential factors: (*i*) advancements in molecular diagnostic methods and novel detection technologies for phytoplasmas^41,42^; (*ii*) a rise in the abundance of leafhopper vectors^31^; (*iii*) the probable existence of yet-to-be-identified highly efficient vectors^22,39,43,44^; and (*iv*) an expansion of the geographic range of leafhopper vectors^23^.

Nonetheless, a significant research gap still exists concerning investigating the potential impacts of climate change on the Nearctic leafhoppers. Our work addresses this gap by examining the potential impact of climate change (abiotic factors) on current and future potential environmental niche similarity and species richness of 14 Nearctic leafhopper species associated with phytoplasmas affecting crops in North America (Table 1; Fig. 1)^34,38–40,43^, employing ENM-based methods. The study is particularly urgent given the scarcity of research on the dynamics of these leafhopper species in North America and the impact of climate change in the short and long term.

**Figure 1.**
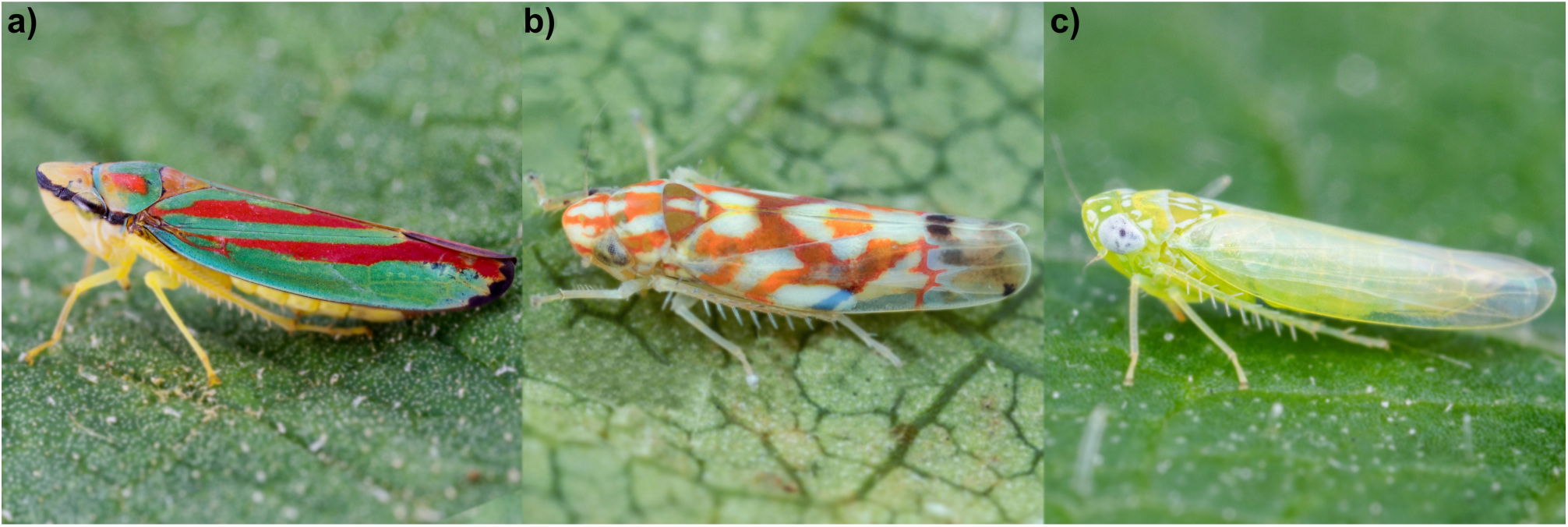
Examples of Nearctic leafhopper genera: a) *Graphocephala* spp., b) *Erythroneura* spp., and c) *Empoasca fabae* (Harris). Joseph Moisan-De-Serres provided pictures from the Ministry of Agriculture, Fisheries and Food – Québec (MAPAQ).

**Table 1.** List of 14 Nearctic leafhopper species studied, their vector status, example crop hosts, and associated phytoplasma diseases.

| Species | §Vector status | Crop hosts | Disease association |
| --- | --- | --- | --- |
| *Amplicephalus inimicus* | Confirmed<sup>45</sup> | Cereals | Aster yellow |
| *Ceratagallia humilis* | Suspected<sup>*43</sup> | Cereals, grass | - |
| *Colladonus geminatus* | Confirmed<sup>46</sup> | Cereals, legumes | Aster yellow |
| *Edwardsiana rosae* | Suspected<sup>*43</sup> | Horticultural, shrubs | - |
| *Empoasca fabae* | Suspected<sup>*43,44</sup> | Field, horticultural, and<br>vegetables | - |
| *Erythroneura comes* | Suspected<sup>#22</sup> | Horticultural | - |
| *Erythroneura vitis* | Suspected<sup>#22</sup> | Horticultural | - |
| *Erythroneura ziczac* | Suspected<sup>*43</sup> | Horticultural | - |
| *Exitianus exitiosus* | Suspected<sup>*43</sup> | Horticultural | - |
| *Graphocephala fennahi* | Suspected<sup>#22</sup> | Shrubs | - |
| *Jikradia olitoria* | Potential/Putative<sup>39</sup> | Horticultural | Grapevine yellows |
| *Macrosteles quadrilineatus* | Confirmed<sup>47,48</sup> | Cereals, field, and<br>vegetables | Aster yellow |
| *Paraphlepsius irroratus* | Confirmed<sup>49</sup> | Horticultural | - |
| *Scaphoideus titanus* | Confirmed<sup>50</sup> | Horticultural | Grapevine yellows |

§The vector status categories indicate: (i) confirmed - the species can acquire and transmit phytoplasmas to crops, (ii) potential - the species can acquire phytoplasmas under laboratory conditions but transmission in the field is not confirmed, and (iii) suspected - phytoplasma sequences have been detected in the leafhopper (\*) or the leafhopper has been found in areas with disease outbreaks but no phytoplasma strains were isolated from the insects (#).

## METHODS

In this study, our objective was to assess environmental similarity and potential leafhopper species richness under current and future climate scenarios using ENM-based methods. The analyses were conducted using the Java version of the machine learning algorithm Maximum Entropy Modeling of Species Geographic Distributions (MaxEnt, v. 3.4.1)^51^, the wallace shiny app (v. 2.0.5)^52^, SDMtoolbox (Pro v. 0.9.1)^53^, ArcGIS Pro (v. 3.1.2), and R packages described further. We chose MaxEnt as our modeling algorithm due to its high statistical performance, mainly when dealing with various species occurrence records datasets^11,54,55^.

### Species selection

We selected 14 species that are confirmed, potentially/putative, or suspected to be vectors of phytoplasmas in North America’s several groups of crops: *Amplicephalus* (formerly *Endria*) *inimicus* (Say), *Ceratagallia* (formerly *Aceratagallia*) *humilis* (Oman), *Colladonus geminatus* (Van Duzee), *Edwardsiana rosae* (Linnaeus), *Empoasca fabae* (Harris), *Erythroneura comes* (Say), *Erythroneura vitis* (Harris), *Erythroneura ziczac* (Walsh), *Exitianus exitiosus* (Uhler), *Graphocephala fennahi* (Young), *Jikradia olitoria* (Say), *Macrosteles quadrilineatus* (formerly *fascifrons*) (Forbes), *Paraphlepsius irroratus* (Say), and *Scaphoideus titanus* (Ball) (Table 1) ^12,22,33,34,39,40,43^. Twelve out of 14 are exclusively present in the Nearctic region. Meanwhile, *Empoasca fabae* extends its occurrence to the Neotropics^56^, while *Scaphoideus titanus*, introduced to Europe in 1958^57^, subsequently spread in this continent due to human trade activities^58^.

### Occurrence records

We obtained species occurrence records (latitude, longitude) for North America from openly available online repositories, including the Global Biodiversity Information Facility (GBIF, accessed on 9 June 2023; https://doi.org/10.15468/dl.h3x8mf), which also includes records from INaturalist platform. We also included recent literature on leafhoppers’ field surveys^22,31,43,44,59^. These records were collected after 1970 to ensure relevance to the environmental variables used in our models (1970-2000) while avoiding historical records that may not accurately reflect current species occurrence sites^60,61^. We removed duplicate records within the same pixel (i.e., the same grid cell) to enhance data quality and eliminate those with incomplete information. We also applied spatial thinning to reduce spatial bias, retaining one record for each 5 km², corresponding to the spatial resolution used in our models. The final number of occurrence records for each species were 300 for *Amplicephalus inimicus*, 33 for *Ceratagallia humilis*, 33 for *Colladonus geminatus*, 66 for *Edwardsiana rosae*, 852 for *Empoasca fabae*, 71 for *Erythroneura comes*, 189 for *Erythroneura vitis*, 110 for *Erythroneura ziczac*, 578 for *Exitianus exitiosus*, 912 for *Graphocephala fennahi*, 2376 for *Jikradia olitoria*, 277 for *Macrosteles quadrilineatus*, 280 for *Paraphlepsius irroratus*, and 94 for *Scaphoideus titanus* (Fig. 2).

**Figure 2.**
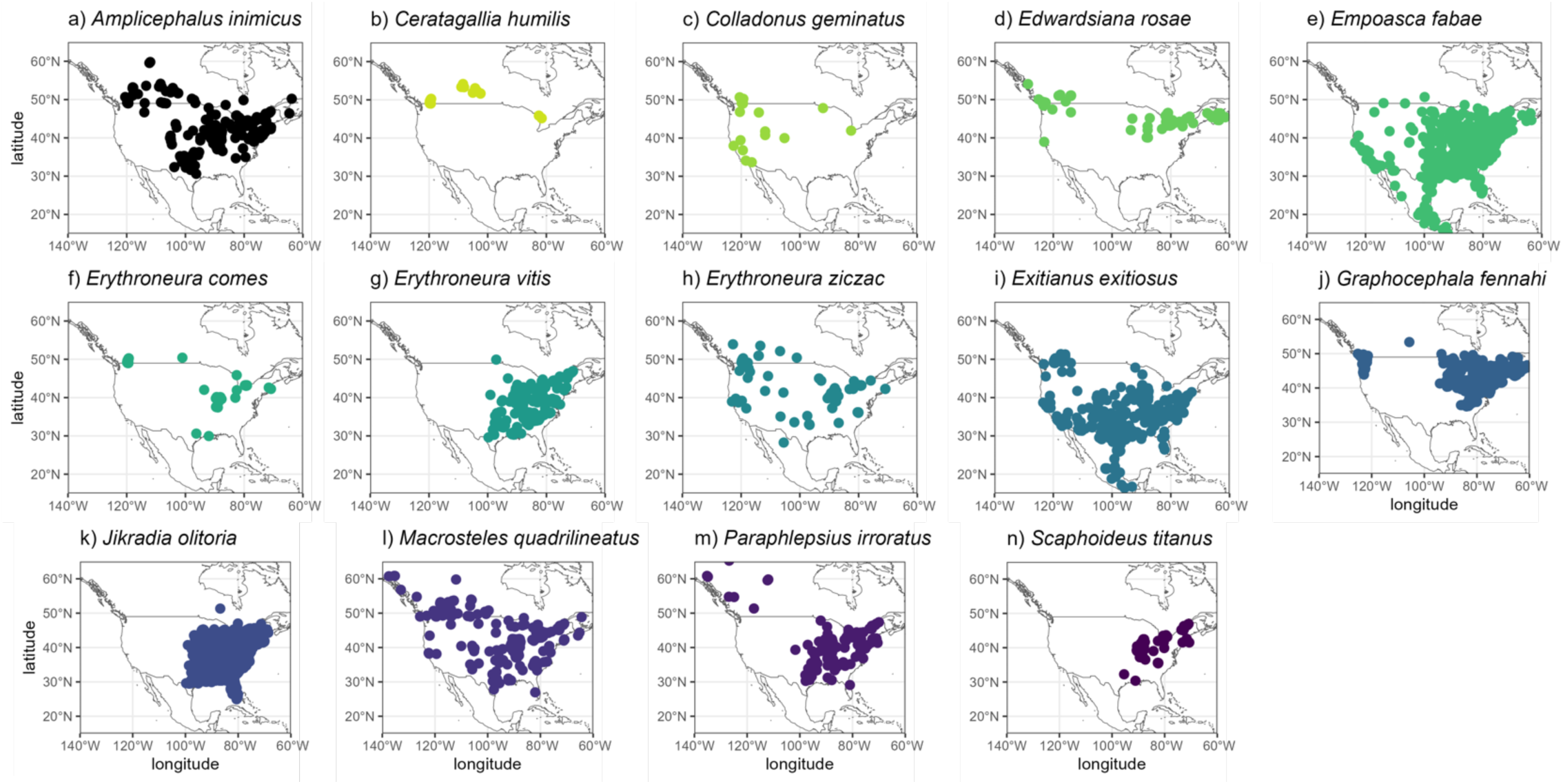
Occurrence records of 14 leafhopper species in North America after removing records with uncertainties and spatial thin (5 km).

### Current environmental variables and climate change scenarios

For our current distribution model, we downloaded 19 world bioclimatic variables (v. 2.1) covering the period of 1970-2000 at a spatial resolution of approximately 5 km²^62^. These variables are derived from monthly temperature and precipitation data, providing insights into annual trends, seasonality, and extreme environmental conditions. Given that these variables can be highly correlated, potentially affecting model performance due to multicollinearity^63^, we conducted a Pearson pair-wise correlation analysis^60^. This analysis allowed us to identify and subsequently remove variables with correlations exceeding a threshold of r ≥ |0.7|, as values beyond this threshold can distort model estimation (Supplementary Table S1)^63^. Thus, five bioclimatic variables were selected: annual mean temperature (bio_1), mean temperature diurnal range (bio_2), temperature annual range (bio_7), annual precipitation (bio_12), and precipitation seasonality (bio_15).

For the climate change scenarios, we used the global circulation model GISS-E2-1-G^64^ under the four shared socioeconomic pathway scenarios, SSP1-2.6, SSP2-4.5, SSP3-7.0, and SSP5-8.5 for the period 2041-2060 at a spatial resolution of 5 km^2^ (available at https://www.worldclim.org/data/cmip6/cmip6_clim2.5m.html).

The GISS-E2-1-G model represents an updated version of the GISS-E2 developed by the National Aeronautics and Space Administration (NASA) Goddard Institute for Space Studies (GISS). This updated model incorporates significant improvements in the representation of climate variability^64^. One notable improvement is the model’s ability to depict the temperature response to increased carbon dioxide. In this iteration, the response is marginally more significant than in previous versions, indicating a more accurate and realistic representation of climate dynamics^64^.

The SSPs, as featured in the Intergovernmental Panel on Climate Change (IPCC) sixth assessment report, offer global narratives that consider responses to climate change. They are associated with various challenges for mitigation and adaptation, ranging from 1 to 5, where 1 represents low, 2 medium, 3 high, 4 low for mitigation and high for adaptation, and 5 high for mitigation and low for adaptation^65^. These SSPs are coupled with global circulation models (GCM) scenarios (previously known as Representative Concentration Pathways – RCP). In our study, we adopted four GCM scenarios: 2.6 (low), 4.5 (intermediate), 7.0 (high), and 8.5 (very high). The SSP1-2.6 scenario limits global warming by 2100 to 2 °C, while the SSP2-4.5, SSP3-7.0, and SSP5-8.5 scenarios project values of 3 °C, 4 °C, and over 4 °C, respectively^65^.

In North America, predictive models indicate significant shifts in the annual mean temperature, demonstrating an increase in maximum values from 28.6 °C to 31 °C, mainly in the southern areas (Supplementary Figure S1). Moreover, the annual precipitation patterns suggest a uniform decrease in the west compared to the east in all scenarios. In contrast, precipitation seasonality is predicted to show the opposite way (Supplementary Figure S1).

We used the raster package in R^66^ to extract bioclimatic variables for each SSP. Subsequently, we created a North American shapefile, and all layers were cropped to align with this shapefile. This cropping procedure was conducted using ArcGIS.

### Model development, selection, and evaluation

MaxEnt is a correlative maximum entropy-based algorithm that estimates species probability of presence or habitat suitability, ranging from 0 (unsuitable) to 1 (highly suitable), using species occurrence records and background data associated with the environmental variables used in the modeling^55^. We used the MaxEnt java (version 3.4.1) and the ENMeval package^67^. We randomly selected 10000 background points to represent environmental conditions across North America for model calibration. We utilized Maxent’s’ clamping’ function to prevent model extrapolations beyond the environmental range. We configured two main sets for model performance and transferability testing: feature class (FC) and regularization multiplier (RM). FC involves mathematical transformations of the environmental variables into predictors: linear (L), quadratic (Q), product (P), and hinge (H), as well as their combinations. RM penalizes model complexity and reduces the number of parameters^68^. In our approach, we tested five FC combinations (L, LQ, H, LQH, LQHP) and RM values ranging from 0 to 5 with a step of 0.5.

We employed two occurrence partition methods, spatial partition block, and random k-fold, to perform cross-validation during model training and testing. The spatial partition method divides localities into four groups based on latitudinal and longitudinal lines, creating spatial rectangles. The random k-fold method randomly assigns localities to one of the ’k’ groups (k = 4). In both approaches, we conducted cross-validation by generating predictions with different training sets and evaluated model performance by projecting outcomes for the omitted sites^68^.

We initially ran 100 models for each species using the feature class (FC) and regularization multiplier (RM) mentioned earlier, combining partition methods and rendering 2800 models. However, we found that the random k-fold method produced better results. The spatial partition block method could not be implemented for some species due to a limited number of localities^52^. Consequently, we retained the random k-fold method for training and testing data. We selected the best model from the Area Under the Receiver Operating Characteristic Curve (AUC) and True Skill Statistics.

The AUC is a standard metric for assessing model performance. It ranges from 0 to 1 and measures the model’s ability to rank positive records higher than negative ones across the suitability range. Values closer to 1 indicate excellent model performance. However, for presence-only models like MaxEnt, interpreting AUC has some limitations, especially when there is a lack of absence data^69^. Additionally, when occurrence records are fewer than 100, AUC may not accurately estimate model complexity^70^. We also used True Skill Statistics (TSS) to address these limitations.

TSS assesses model performance by considering sensitivity and specificity at a binary threshold (unsuitable/suitable). TSS values range from -1 to 1, with values closer to one indicating excellent sensitivity and specificity. High TSS (> 0.5) values mean a high proportion of real presences, and current absences are correctly predicted. Conversely, values at or below zero suggest errors in predicting presence (commission) or predicting absence (omission)^60,71^.

### Projections

We transformed asci (cloglog output) maps into raster files using ArcGIS. These maps were then classified into binary outputs, unsuitable and suitable, for each species based on the maximum test sensitivity plus specificity cloglog threshold (MaxTSS values in Table 1). Sensitivity and specificity represent the proportions of correctly predicted presences and absences, respectively. Subsequently, we overlaid the binary maps to calculate species richness under current and future scenarios. We used the ’biodiversity measurement’ function available in SDMtoolbox. The final output is a map showing the sum of unique species per pixel, representing the total number of species in each grid cell.

### Environmental similarity

We assessed environmental similarity using the final species occurrence records and the five environmental variables employed in our models under current and four future scenarios. We conducted a principal component analysis (PCA) to reduce the original environmental space’s dimensionality. Subsequently, we calculated species occurrence density along the first two components of the PCA and performed similarity tests.

In the similarity test, both species’ environments were randomly shifted within the background extent, and ’null’ overlaps were calculated from these simulations. If the observed overlap was greater than 95% of the simulated overlaps (*p*-value < 0.05), the test indicates that the two species’ environments were more similar than expected by chance (a one-tailed test). We conducted tests by pairing two leafhopper species at each iteration until all 91 possible combinations among the 14 species were evaluated for each climate scenario. Subsequently, we created a matrix containing the similarity *p*-values (supplementary Table S2).

The similarity *p*-values matrix was further transformed into a binary matrix, where ’0’ represents random, and ’1’ indicates species with environment similarity (supplementary Table S2). Using network analysis, we identified species with higher and lower similarity. For each network scenario, we calculated the network density. Density is the proportion of observed ties or connections to a network’s maximum and possible relations. Thus, density is a ratio that can range from 0 to 1. The closer to 1 the density is, the more interconnected the network^72^. These analyses and tests were performed using the wallace interface to compute the tests, the packages ade4^73^, igraph^74^, ecospat^75^, and Corel Painter (Essential 7, ON, CAD) for data illustration.

## RESULTS

### Models performance and variables contribution

Our predictive models demonstrated high discriminant performance, as evidenced by AUC values ranging from 0.883 to 0.980 and TSS values spanning from 0.629 to 0.864 (Table 2). Our relative contribution analysis highlighted that temperature-related variables substantially shaped the models for most of the studied species (Fig. 3). However, some exceptions were observed for *Erythroneura comes*, *Erythroneura vitis*, *Graphocephala fennahi*, and *Jikradia olitoria*, where precipitation-related variable had a more substantial influence on the models. Interestingly, in the case of *Ceratagallia humilis* (Temperature: 51.11%; Precipitation: 48.89%) and *Scaphoideus titanus* (Temperature: 50.03%; Precipitation: 49.97%), both sets of variables contributed almost equally to the model (Fig. 3).

**Figure 3.**
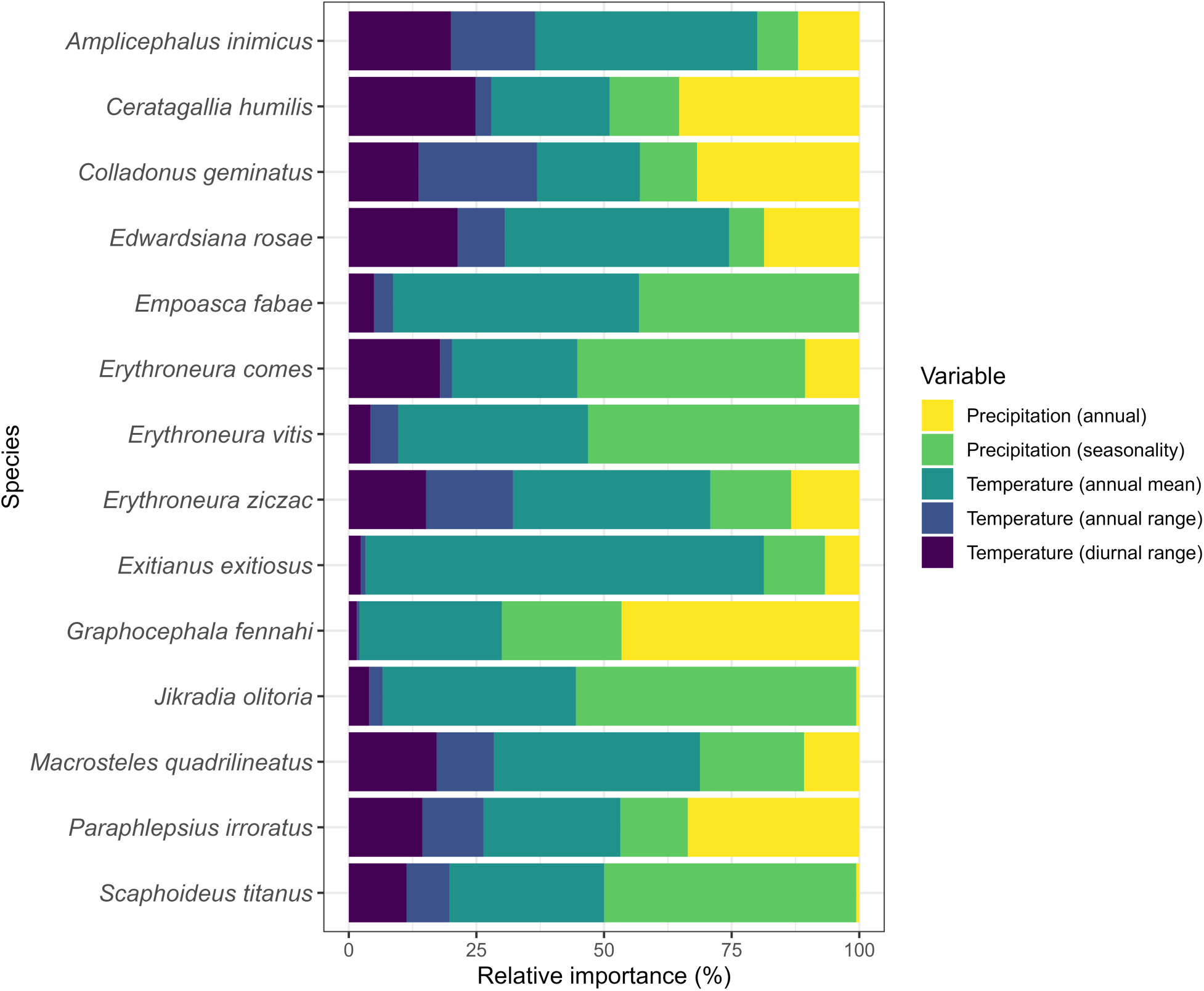
The relative importance of the environmental variables for the current potential distribution models for 14 leafhopper species in North America.

**Table 2.** Selected model setup and performance metrics for 14 leafhopper species in North America. Feature class (FC) and regularization multiplier (RC) indicated the model setup in MaxEnt. Area under the operational curve (AUC) and true skill statistics (TSS) indicate model performance. MaxTSS refers to the probability threshold for binary conversion of the maps (unsuitable and suitable). SD = standard deviation (n = 4).

| Species | FC | RM | AUC (SD) | TSS (SD) | MaxTSS |
| --- | --- | --- | --- | --- | --- |
| *Amplicephalus inimicus* | LQH | 0.5 | 0.928 (0.012) | 0.723 (0.030) | 0.166 |
| *Ceratagallia humilis* | H | 0.5 | 0.980 (0.012) | 0.828 (0.009) | 0.066 |
| *Colladonus geminatus* | LQH | 1.5 | 0.918 (0.051) | 0.700 (0.080) | 0.057 |
| *Edwardsiana rosae* | LQH | 4.0 | 0.948 (0.018) | 0.766 (0.081) | 0.373 |
| *Empoasca fabae* | L | 0.5 | 0.883 (0.011) | 0.706 (0.028) | 0.341 |
| *Erythroneura comes* | LQH | 4.0 | 0.974 (0.013) | 0.852 (0.077) | 0.144 |
| *Erythroneura vitis* | LQ | 4.0 | 0.954 (0.006) | 0.832 (0.017) | 0.379 |
| *Erythroneura ziczac* | LQH | 1.5 | 0.939 (0.018) | 0.738 (0.069) | 0.222 |
| *Exitianus exitiosus* | LQ | 1.0 | 0.916 (0.007) | 0.698 (0.005) | 0.218 |
| *Graphocephala fennahi* | H | 4.0 | 0.939 (0.004) | 0.840 (0.003) | 0.247 |
| *Jikradia olitoria* | LQ | 0.5 | 0.888 (0.004) | 0.816 (0.004) | 0.219 |
| *Macrosteles quadrilineatus* | LQHP | 0.5 | 0.895 (0.017) | 0.629 (0.051) | 0.260 |
| *Paraphlepsius irroratus* | LQHP | 0.5 | 0.953 (0.011) | 0.810 (0.054) | 0.264 |
| *Scaphoideus titanus* | LQ | 0.5 | 0.979 (0.006) | 0.864 (0.033) | 0.050 |

### Current species richness

The richness map obtained in this study reveals that the eastern part of North America, encompassing southern Ontario and Québec—identified as an anticipated high richness region for Nearctic leafhoppers^13,17^—exhibits the potential for the highest species richness (Fig. 4). Another high richness area was observed in the southwestern of the United States. These regions typically feature a temperate climate with elevated humidity levels. In contrast, richness decreases in drier (e.g., the grasslands of Canada and the U.S., which include per-arid, arid, and semi-arid zones) and colder areas (e.g., Great Plains). Consequently, northern regions of Canada and the U.S. show a lower richness (Fig. 4).

**Figure 4.**
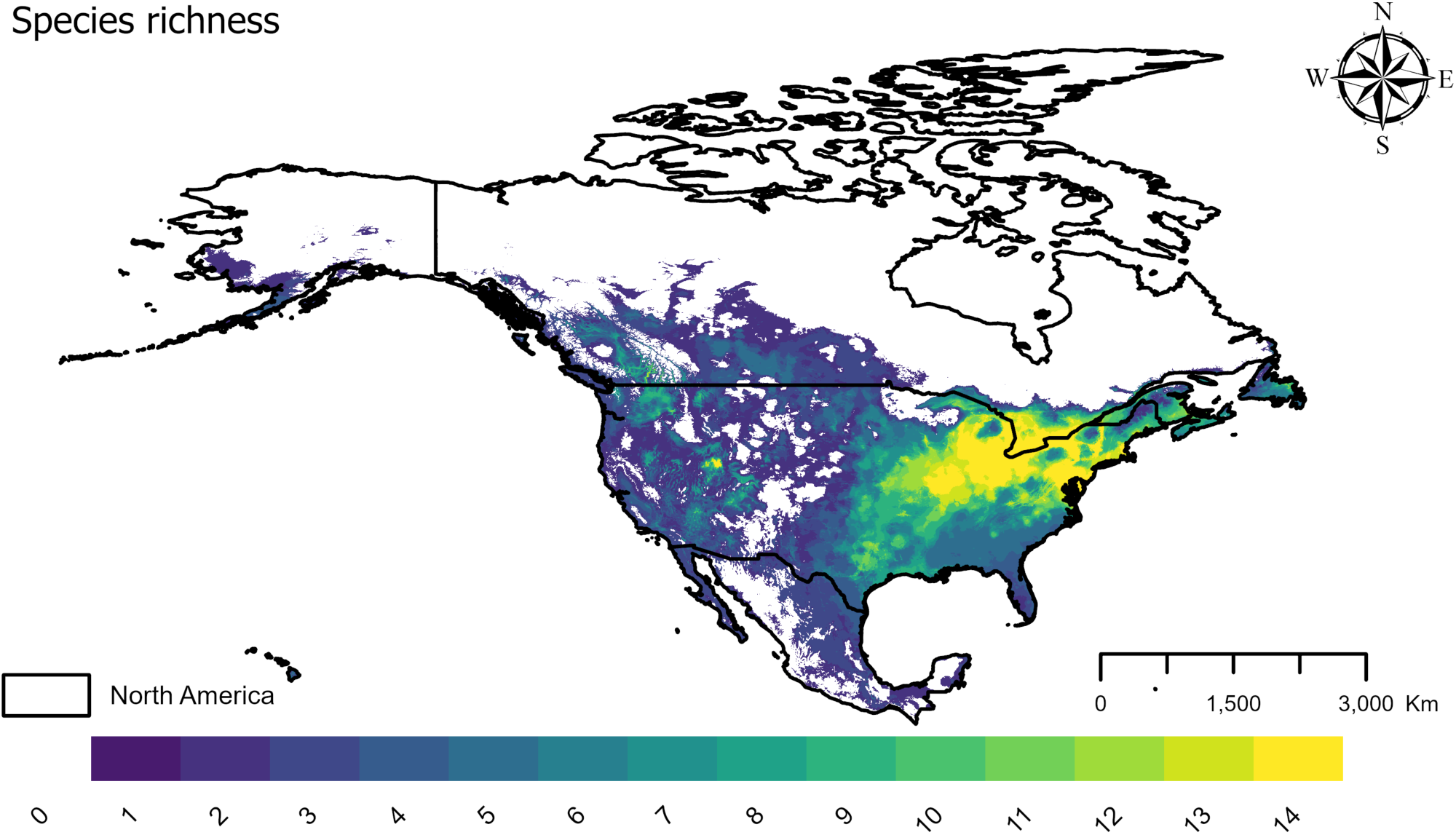
Estimated current leafhopper species richness based on 14 species potential distribution models in North America.

### Future projections

To predict the impact of climate change on leafhopper richness, future scenarios SSP1-2.6 (Fig. 5a), 2-4.5 (Fig. 5b), 3-7.0 (Fig. 5c), and 5-8.5 (Fig. 5d) were evaluated, finding that under the four scenarios, suitable areas for species richness will tend to increase (Table 3). The regions previously identified with the highest species richness, hosting all 14 species, are projected to retain similar patterns in all four future scenarios (Fig. 5). We observed a considerable increase in this richness, gradually extending towards the east (Fig. 5). A new high richness area emerged in the southern parts of Alberta and British Columbia, Canada (Fig. 5). Additionally, an expansion in richness is evident in the southeast and northeast parts of the continent. In line with current conditions, our models predict lower species richness in arid zones compared to regions with higher humidity (Fig. 5). Altogether, our results indicate that, under the four climate change scenarios investigated, northern Canada is expected to become more hospitable to a more significant number of species than under the current scenario.

**Figure 5.**
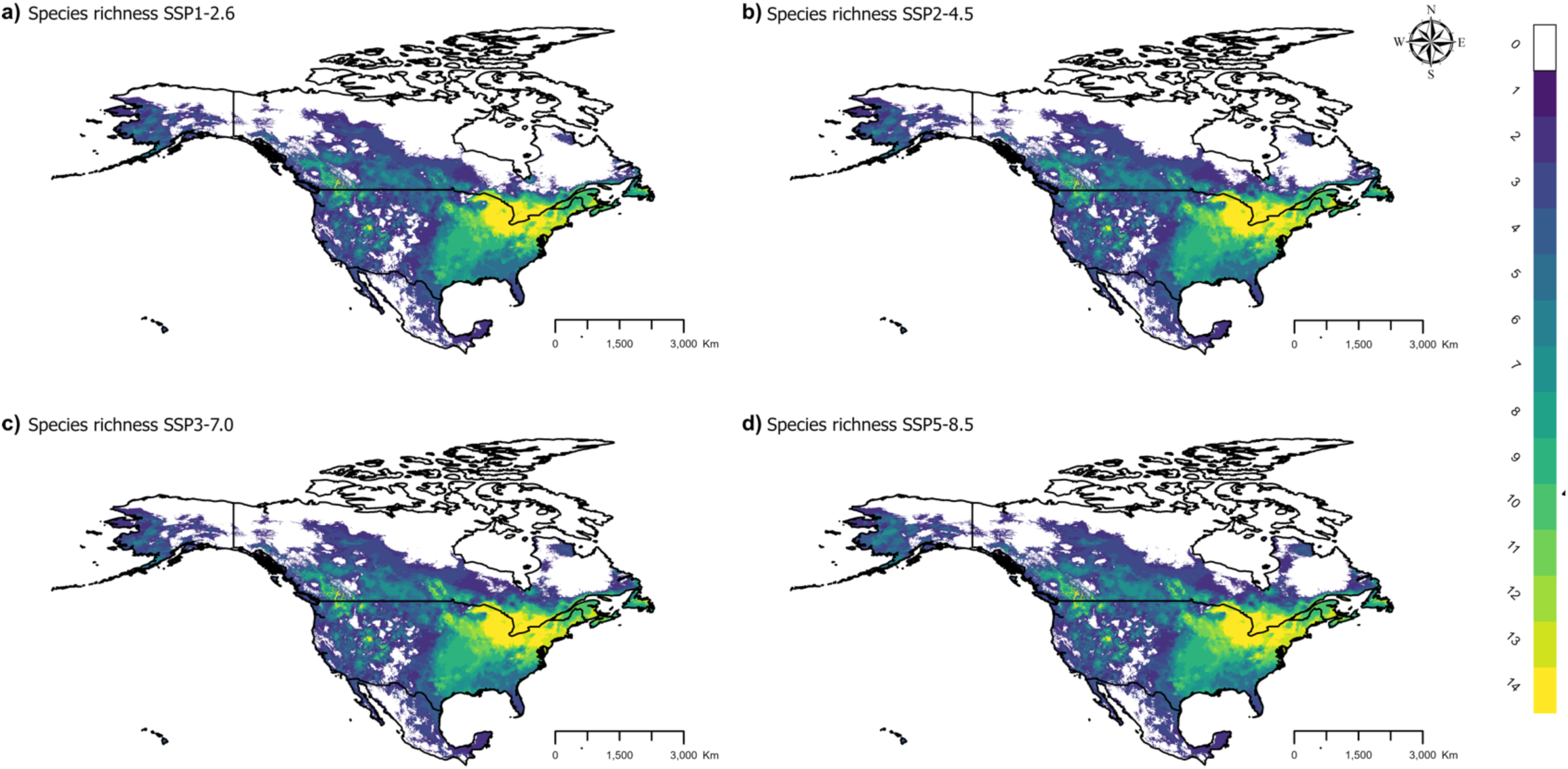
Estimated leafhopper species richness based on 14 species potential distribution in North America under four shared socioeconomic pathway scenarios [a) SSP1-2.6, b) SSP2-4.5, c) SSP3-7.0, and d) SSP5-8.5) for 2041-2060.

**Table 3.**
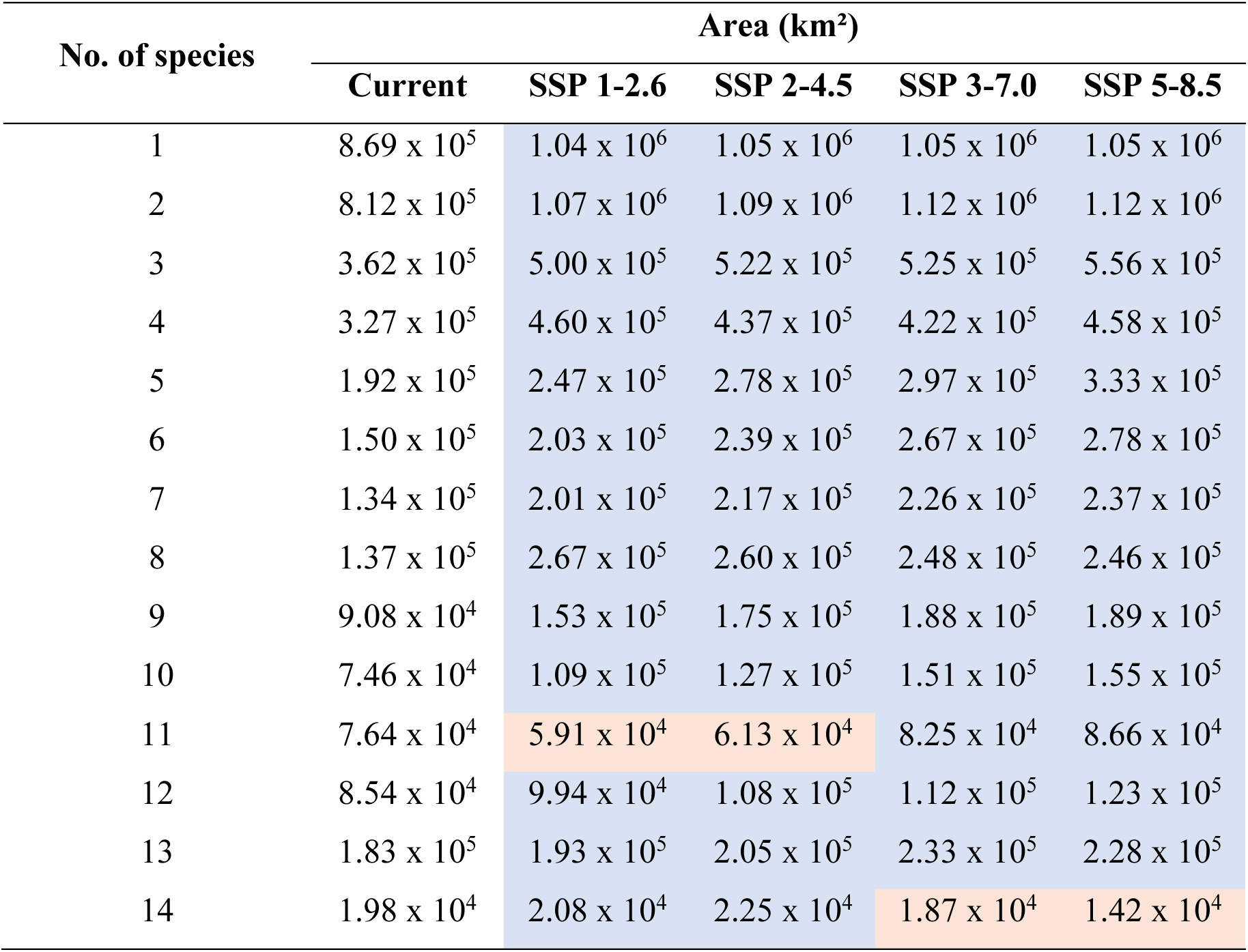
Estimated area extent for species richness (number of species) in North America under current and four shared socioeconomic pathway scenarios (SSP1-2.6, SSP2-4.5, SSP3-7.0, and SSP5-8.5) for 2041-2060. Blue indicates an increase in suitable areas, and red indicates a decrease compared to the current scenario.

| No. of species | Area (km<sup>2</sup>) |  |  |  |  |
| --- | --- | --- | --- | --- | --- |
|  | Current | SSP 1-2.6 | SSP 2-4.5 | SSP 3-7.0 | SSP 5-8.5 |
| 1 | 8.69 x 10<sup>5</sup> | 1.04 x 10<sup>6</sup> | 1.05 x 10<sup>6</sup> | 1.05 x 10<sup>6</sup> | 1.05 x 10<sup>6</sup> |
| 2 | 8.12 x 10<sup>5</sup> | 1.07 x 10<sup>6</sup> | 1.09 x 10<sup>6</sup> | 1.12 x 10<sup>6</sup> | 1.12 x 10<sup>6</sup> |
| 3 | 3.62 x 10<sup>5</sup> | 5.00 x 10<sup>5</sup> | 5.22 x 10<sup>5</sup> | 5.25 x 10<sup>5</sup> | 5.56 x 10<sup>5</sup> |
| 4 | 3.27 x 10<sup>5</sup> | 4.60 x 10<sup>5</sup> | 4.37 x 10<sup>5</sup> | 4.22 x 10<sup>5</sup> | 4.58 x 10<sup>5</sup> |
| 5 | 1.92 x 10<sup>5</sup> | 2.47 x 10<sup>5</sup> | 2.78 x 10<sup>5</sup> | 2.97 x 10<sup>5</sup> | 3.33 x 10<sup>5</sup> |
| 6 | 1.50 x 10<sup>5</sup> | 2.03 x 10<sup>5</sup> | 2.39 x 10<sup>5</sup> | 2.67 x 10<sup>5</sup> | 2.78 x 10<sup>5</sup> |
| 7 | 1.34 x 10<sup>5</sup> | 2.01 x 10<sup>5</sup> | 2.17 x 10<sup>5</sup> | 2.26 x 10<sup>5</sup> | 2.37 x 10<sup>5</sup> |
| 8 | 1.37 x 10<sup>5</sup> | 2.67 x 10<sup>5</sup> | 2.60 x 10<sup>5</sup> | 2.48 x 10<sup>5</sup> | 2.46 x 10<sup>5</sup> |
| 9 | 9.08 x 10<sup>4</sup> | 1.53 x 10<sup>5</sup> | 1.75 x 10<sup>5</sup> | 1.88 x 10<sup>5</sup> | 1.89 x 10<sup>5</sup> |
| 10 | 7.46 x 10<sup>4</sup> | 1.09 x 10<sup>5</sup> | 1.27 x 10<sup>5</sup> | 1.51 x 10<sup>5</sup> | 1.55 x 10<sup>5</sup> |
| 11 | 7.64 x 10<sup>4</sup> | 5.91 x 10<sup>4</sup> | 6.13 x 10<sup>4</sup> | 8.25 x 10<sup>4</sup> | 8.66 x 10<sup>4</sup> |
| 12 | 8.54 x 10<sup>4</sup> | 9.94 x 10<sup>4</sup> | 1.08 x 10<sup>5</sup> | 1.12 x 10<sup>5</sup> | 1.23 x 10<sup>5</sup> |
| 13 | 1.83 x 10<sup>5</sup> | 1.93 x 10<sup>5</sup> | 2.05 x 10<sup>5</sup> | 2.33 x 10<sup>5</sup> | 2.28 x 10<sup>5</sup> |
| 14 | 1.98 x 10<sup>4</sup> | 2.08 x 10<sup>4</sup> | 2.25 x 10<sup>4</sup> | 1.87 x 10<sup>4</sup> | 1.42 x 10<sup>4</sup> |

### Environmental similarity

Our network analysis revealed that most leafhopper species present a similar environmental niche based on the five variables used in the modelling with a density value of 0.467 (Fig. 6). Notably, *Colladonus geminatus*, *Edwardsiana rosae,* and *Exitianus exitiosus* showed a remarkable environmental similarity with other 4 and 3 species, respectively, indicating a lesser environmental similarity. In contrast, *Amplicephalus inimicus*, *Empoasca fabae*, *Erythroneura* spp., *Macrosteles quadrilineatus*, *Paraphlepsius irroratus*, and *Scaphoideus titanus* exhibited a higher degree of environmental similarity with most of the species in our study, suggesting a higher environmental niche similarity (Fig. 6). These findings underscore the existence of a distinct environmental niche similarity between the 14 leafhopper species studied here.

**Figure 6.**
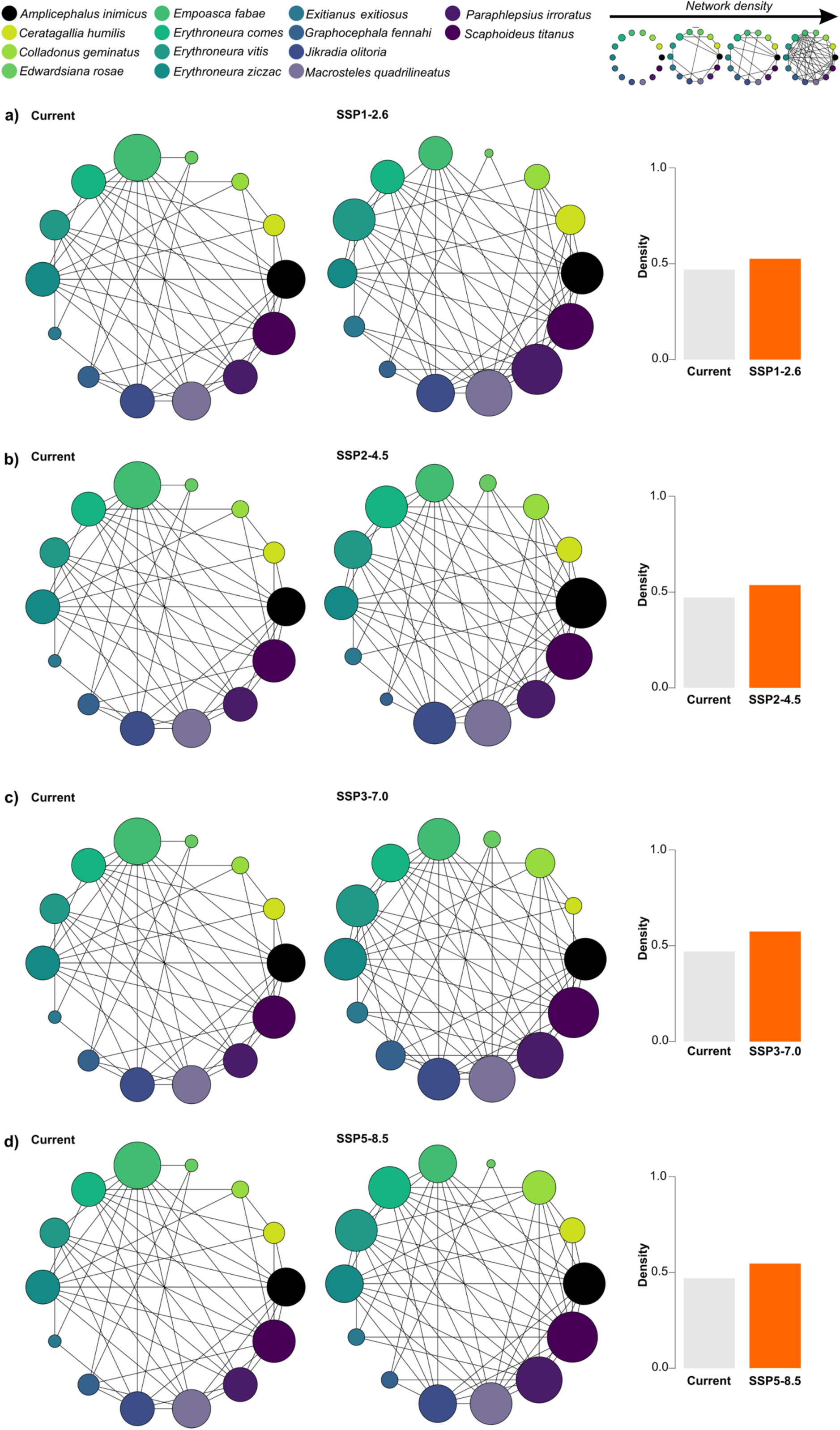
A comparison of environmental similarity networks and their density among 14 leafhopper species in North America under current and four future scenarios: a) SSP1-2.6, b) SSP2-4.5, c) SSP3-7.0, and d) SSP5-8.5 for the period 2041-2060. The columns, represented in grey for the current scenario and orange for the four alternative scenarios, indicate the network density values. This metric reflects the proportion of observed connections relative to the network’s maximum potential connections. Density is a ratio that ranges from 0 to 1, with higher values indicating a more interconnected network. The size of circles within the network graph represents the number of interactions between species, meaning larger circles signify more significant similarities among species.

Under four future scenarios, the environmental similarity tended to increase between leafhopper species suggesting more interconnected networks as indicated by the increase of density values from 0.467 under the current scenario to 0.523 under SSP1-2.6 (Fig. 6a), 0.533 under SSP2-4.5 (Fig. 6b), 0.571under SSP3-7.0 (Fig. 6c), and 0.542 under SSP5-8.5 (Fig. 6d). In all scenarios, leafhopper species tended to present environmental similarity, including those with less degree of environmental niche similarity under the current scenario like *Colladonus geminatus* and *Edwardsiana rosae*. An exception was registered for *Exitianus exitiosus*, which only shares an environmental similarity with a maximum of three species. Overall, our results underscore the potential of leafhopper species to occupy a more similar environment niche under a warming world scenario.

## DISCUSSION

Our study examined the environmental niche similarity and distribution of 14 Nearctic leafhopper species across North America under current conditions and four future climate scenarios. We found that 11 out of 14 leafhopper species share a similar environmental niche with more species. Meanwhile, *Colladonus geminatus*, *Edwardsiana rosae*, and *Exitianus exitiosus* show less similarity in the environmental niche. Interestingly, a similar trend appears to grow across climate change scenarios. Our prediction points to warm and humid regions in North America as potential high-richness areas, with fewer species expected to occur in arid and colder areas. Under all climate scenarios, we observed an expansion of the suitable regions and species overlap, especially in the northern area of the continent. Our findings suggest a positive link between future climates and the potential occurrence and environmental niche similarity of 14 North America’s Nearctic leafhopper species with some degree of association with phytoplasmas (Table 1).

According to our models, temperature significantly influences the distribution of most leafhopper species investigated in this study. In the Nearctic region, temperature is the primary factor driving leafhopper distribution. Historical periods of high temperatures are thought to trigger large-scale dispersal of leafhopper species into northern areas^16,19^. During winter, some leafhopper species undergo temperature-related diapause in their egg, nymph, or adult stages^14,19,76^, while migratory species such as *Empoasca fabae* overwinter in southern regions^23^. As temperatures rise in early spring, leafhopper populations start to increase from both diapausing stages and the arrival of migratory species due to continental-scale winds^24,25^, with populations peaking in warmer months^22,23,31^. Leafhopper populations in the Nearctic region exhibit shorter life cycles in warmer conditions^21^, resulting in more yearly generations than colder ones^23,76^. Yet, precipitation also plays a role in species occurrence, particularly noted in our models for *Erythroneura comes*, *Erythroneura vitis*, *Graphocephala fennahi*, and *Jikradia olitoria*. Heavy precipitation, especially during early nymphal stages, affects the mobility of leafhoppers and can cause species mortality^13^. In addition, high humidity indirectly impacts species’ performance by affecting their plant hosts, as noted for *Erythroneura* spp., which tends to increase populations due to weakened host plants due to moisture^76^. Upon conducting an environmental niche analysis, we show substantial environmental similarity among most leafhopper species. As previously mentioned, historical dispersion patterns of leafhoppers indicate movements from regions characterized by higher diversity and warmer climates to regions with lower diversity and cooler climates, such as between the Neotropical and Nearctic regions^16,19,77^. Consequently, approximately 90% of leafhopper genera exhibit endemism within their respective zoogeographical regions, demonstrating a marked inclination toward host specificity and limited dispersal rates^16^. This endemism potentially elucidates the environmental similarity observed among the species under study, given that the majority, 12 out of 14, exclusively inhabit the Nearctic region and only *Empoasca fabae* is known to not overwinter in Canada^13,14,76,78^.

Recent surveys suggest potential overlap and environmental niche similarity among some Nearctic leafhopper species. These observations are based on initial evidence of increased species richness in specific regions. For instance, Saguez *et al.*^59^ documented 17,496 leafhopper specimens across Canadian vineyards (British Columbia, Québec, and Ontario), spanning 54 genera and 110 species. Notably, the genera *Empoasca*, *Erythroneura*, and *Macrosteles* were found to be widespread. Plante *et al.*^22^ captured 33,007 leafhopper specimens from Quebec strawberry fields, representing 61 genera and 118 species, with *Empoasca*, *Macrosteles*, and *Hebata* prevalent. However, it is crucial to acknowledge that these studies employed different sampling methods, targeted distinct sites and hosts, and were conducted in separate years. Therefore, caution is necessary when interpreting this initial evidence. Further studies with standardized methodologies are needed to confirm this potential trend of species overlap.

In contrast, three species, *Colladonus geminatus*, *Edwardsiana rosae*, and *Exitianus exitiosus,* displayed a reduced level of niche similarity, suggesting a more specialized environmental niche within regions characterized by lower species richness. Intriguingly, two species, *Colladonus geminatus* and *Exitianus exitiosus*, exhibit distribution and abundance within grassland habitats^13,79^, which are recognized for harboring reduced species diversity compared to humid environments^13,16^. Our assessment of species richness has uncovered a geographical trend for the species evaluated: areas with higher humidity and moisture generally support a greater variety of species, with numbers dwindling in drier and colder regions. This prediction aligns with prior observations in southern Ontario/Québec^13^ and New Hampshire^17^, areas known for high biodiversity. These findings suggest a complex relationship between environmental conditions and the resulting distribution of leafhopper species.

Under the four climate change scenarios studied, our network analysis revealed a notable rise in species similarity, hinting at a likely increase of species overlap within their suitable habitats in forthcoming scenarios^8^. Some initial indicators emphasize the impact of rising temperatures on the species associated studied, leading to larger distribution areas for these species. For example, the northward shift of the migratory *Empoasca fabae* in North America has sped up over the last 62 years, a change linked to warmer temperatures^23^. This earlier arrival has notably intensified infestations, potentially favoring this species in the North American region^23^. A recent survey conducted in Québec’s strawberry fields identified 11 leafhopper species for the first time in the province. Ten of these species were the farthest observed towards the east in Canada, marking a significant expansion^22^. This survey also noted a considerable increase in leafhopper species composition compared to data from 14 years ago in vineyards^59^ and 10 years ago in blueberry fields^44^, with 53 new species registered in 2021 and 2022^22^. Given our predictive models’ observed range expansion trends, these findings might suggest a growing overlap among Nearctic leafhopper species in North America. However, as previously mentioned, the lack of standardized studies across existing surveys limits comparison. This underscores the need for further research with consistent methodologies to test this hypothesis of widespread Nearctic leafhopper species overlap.

We expect a significant expansion of suitable areas for various leafhopper species, especially in northern areas^3,4^. This trend mirrors similar trends observed in other groups of crop pests, moving toward northern regions driven primarily by global warming^4^. This shift results in broader ranges for species distribution and overwintering zones^3,4,80^. Additionally, multiple lines of evidence support that most species in the Nearctic region are likely to adjust their geographic distribution to occupy areas that meet their specific needs^1,6,8,81^. This adaptability stems from the ability of Nearctic species to cope with seasonal temperatures and extreme variations, survive cold winters, and succeed in hot summers, thus aligning their life cycles with these environmental conditions^6^.

In contrast to Nearctic leafhoppers, Neotropical species appear less adaptable to warm climates due to their specialization within a narrower temperature range and increased sensitivity to precipitation changes. For example, *Macrosteles quadrilineatus* can survive 18 days at 0 °C, but reproduction ceases. Similarly, at 35°C, survival remains at 18 days, but reproduction is still possible^21^. This highlights the contrasting thermal tolerance compared to *Dalbulus maidis* (DeLong & Wolcott), a Neotropical leafhopper, with a developmental temperature range between 20 °C and 30 °C ^82^. A warming climate could significantly reduce suitable habitats for *Dalbulus maidis* in its native South American range as temperatures approach their upper limit^9^.

Our study investigated the impact of abiotic factors on current and future potential environmental niche similarity and species richness of 14 North American leafhopper potential vectors of phytoplasmas. However, they do not account for biotic variables such as natural enemies (parasitoids and predators) and the availability of hosts, the seasonal effect on area suitability, or the effect of climate change on continental wind patterns. These factors, collectively, act as a biological filter, influencing the actual niche of the species studied^19,60^. Moreover, many Nearctic leafhoppers do not inhabit their entire potential range^17,19^. Therefore, our analyses should be viewed through the lens of the environmental niche concept, focusing primarily on the climatic factors limiting species distribution. This approach often extends beyond the realized niche^60^.

Our study is also limited by the lack of ‘historical’ records (pre-1970) that might still reflect the current species occurrence (e.g., *Scaphoideus titanus*). We excluded these records to avoid (i) a temporal bias in our dataset, following established guidelines^60,61^, and (ii) a mismatch with the environmental variables timeline^62^. Additionally, existing literature for some species only provides presence/absence data without geographic coordinates (e.g.,^83^). Furthermore, distinguishing between taxa in species complexes can be challenging (e.g., *Ceratagallia humilis*), making it difficult to determine true absences versus those due to nomenclature issues^18^. Finally, as previously mentioned, the actual distribution of many Nearctic leafhoppers remains unknown or understudied. Despite these limitations, this is the first study to explore the potential richness and environmental similarity of Nearctic leafhopper species across North America’s current and future climate scenarios. Overcoming these limitations, through further studies expanding species occurrence as seen with other taxa such as bees^84^, will be crucial for improving future species distribution predictions.

## IMPLICATIONS FOR SUSTAINABLE AGRICULTURE

Our study highlights significant challenges in managing Nearctic leafhoppers, particularly their continued presence in current habitats and their potential movement towards northern regions in North America. Although insecticide resistance in Nearctic leafhopper species has not been studied in depth, new evidence suggests that the heavy reliance on insecticides for leafhopper control in Canada could be becoming less effective at managing populations^22^. Moreover, the early arrival of migratory leafhopper populations exacerbates plant damage^23^, potentially leading to increased insecticide usage and increasing the chances of developing insecticide resistance^22,85^.

Additionally, the potential resilience of Nearctic leafhopper species under warmer climates could amplify the spread of diseases they transmit. It is well-known that temperature fluctuations moderate phytoplasma multiplication within leafhoppers^21^, and higher temperatures accelerate phytoplasma cycles within plants^86–88^, increasing the chances of acquisition or transmission during vector feeding activities. Given that species such as *Macrosteles quadrilineatus* have already tested positive for phytoplasma strains that affect multiple hosts^38,43^, another anticipated effect of climate change might be the expansion of host ranges for these phytoplasmas, previously limited by the temperature and food sources available to leafhoppers, as previously reported by our group happening with the BbSP phytoplasma strain moving from blueberries^41^ to lingonberry^86^ in Québec, Canada.

The changes in climate patterns will lead to increased Nearctic leafhopper activity, migration, and interaction, which will negatively affect plant health. These results call for a complete rethink of our control and integrated pest management strategies to meet these new challenges. Little is done in North America to manage specifically Nearctic leafhoppers, however, in Brazil after the introduction of Bt-corn, leafhopper species like *Dalbulus maidis* has become a great threat^87^. For example, the use of parasitoids has been explored as a biocontrol strategy^88^, while more recently the use of RNAi technology has shown promising results^89^. Our study emphasizes the need for further research to discover effective products to manage Nearctic leafhopper populations like RNAi, genome edited crops less attractive to these pests, and tailored biocontrol agents. Additionally, we should explore new methods of biological control and the potential of biotechnology and emerging technologies to halt the spread of these pests and the diseases they carry without harming the environment.

## AUTHOR CONTRIBUTION

Conceptualization: A.A.S. and E.P.L.; Data analysis and presentation: A.A.S.; Writing full draft, review, and editing: A.A.S., J.J., and E.P.L.; Supervision: E.P.L.

## DATA AVAILABILITY

The dataset and resources used in this study are openly available on websites upon user registration: GBIF (https://www.gbif.org/), INaturalist (https://www.inaturalist.org/), MaxEnt java (https://biodiversityinformatics.amnh.org/open_source/maxent/), Wallace app (https://wallaceecomod.github.io/), WorldClim climate data (https://www.worldclim.org/data/index.html). Supplementary tables and figures can be found in Zenodo following this doi: https://doi.org/10.5281/zenodo.10371910.

## ACKNOWLEDGMENTS

We extend our gratitude to M.Sc. Joseph Moisan-De-Serres (Ministry of Agriculture, Fisheries and Food, Québec) for the leafhoppers pictures (figure 1) and Dr. Leonardo Turchen (Carleton University, Ottawa) for valuable assistance in preparing figures 1, 5, and 6. We acknowledge the two anonymous reviewers for their helpful comments to improve the manuscript. Additionally, we express our appreciation to the developers of the free database and software resources that were instrumental in facilitating the execution of this study.

## REFERENCES

1. Pecl, G. T. et al. Biodiversity redistribution under climate change: Impacts on ecosystems and human well-being. Science 355, eaai9214 (2017).

2. Kharouba, H. M., Lewthwaite, J. M. M., Guralnick, R., Kerr, J. T. & Vellend, M. Using insect natural history collections to study global change impacts: challenges and opportunities. Phil. Trans. R. Soc. B 374, 20170405 (2019).

3. Bebber, D. P., Ramotowski, M. A. T. & Gurr, S. J. Crop pests and pathogens move polewards in a warming world. Nature Clim. Change 3, 985–988 (2013).

4. Bebber, D. P., Holmes, T. & Gurr, S. J. The global spread of crop pests and pathogens. Glob. Ecol. Biogeogr. 23, 1398–1407 (2014).

5. Fenn-Moltu, G., et al. Global flows of insect transport and establishment: The role of biogeography, trade and regulations. Divers. Distrib. 29, 1478–1491 (2023).

6. Harvey, J. A., Heinen, R., Gols, R. & Thakur, M. P. Climate change-mediated temperature extremes and insects: From outbreaks to breakdowns. Glob. Change Biol. 26, 6685–6701 (2020).

7. Lehmann, P. et al. Complex responses of global insect pests to climate warming. Front. Ecol. Environ. 18, 141–150 (2020).

8. Araújo, M. B. et al. Heat freezes niche evolution. Ecol. Lett. 16, 1206–1219 (2013).

9. Santana, P. A., Kumar, L., Da Silva, R. S., Pereira, J. L. & Picanço, M. C. Assessing the impact of climate change on the worldwide distribution of *Dalbulus maidis* (DeLong) using MaxEnt. Pest. Manag. Sci. 75, 2706–2715 (2019).

10. Andrew, N. R. et al. Assessing insect responses to climate change: What are we testing for? Where should we be heading? PeerJ 1, e11 (2013).

11. Feng, X. et al. A checklist for maximizing reproducibility of ecological niche models. Nat. Ecol. Evol. 3, 1382–1395 (2019).

12. Weintraub, P. G. & Beanland, L. Insect vectors of phytoplasmas. Annu. Rev. Entomol. 51, 91– 111 (2006).

13. Beirne, B. P. Leafhoppers (Homoptera: Cicadellidae) of Canada and Alaska. Can. Entomol. 88, 1–180 (1956).

14. DeLong, D. M. The leafhoppers, or Cicadellidae, of Illinois (Eurymelinae Balcluthinae). Bull. Ill. Nat. Hist. Surv. 24, 97–376 (1948).

15. Bartlett, C. R. et al. The diversity of the true hoppers (Hemiptera: Auchenorrhyncha) 501-590 (ed. Foottit, R. G. & Adler, P. H.) (John Wiley & Sons, Inc, 2017).

16. Nielson, M. W. & Knight, W. J. Distributional patterns and possible origin of leafhoppers (Homoptera, Cicadellidae). Rev. Bras. Zool. 17, 81–156 (2000).

17. Chandler, D. S. & Hamilton, K. G. A. Biodiversity and Ecology of the Leafhoppers (Hemiptera: Cicadellidae) of New Hampshire. Trans. Am. Entomol. Soc. 143, 773–971 (2017).

18. Hamilton, K. G. A. The species of the North American leafhoppers *Ceratagallia* Kirkaldy and *Aceratagallia* Kirkaldy (Rhynchota: Homoptera: Cicadellidae). Can. Entomol. 130, 427–490 (1998).

19. Hamilton, K. G. A. Leafhoppers (Homoptera: Cicadellidae) of the Yukon: Dispersal and Endemism 337-375 (ed. Danks, H. V. & Downes, J. A.) (University of Chicago, 1997).

20. Pinedo-Escatel, J. A., et al. Biogeographical evaluation and conservation assessment of arboreal leafhoppers in the Mexican Transition Zone biodiversity hotspot. Divers. Distrib. 27, 1051– 1065 (2021).

21. Bahar, M. H., Wist, T. J., Bekkaoui, D. R., Hegedus, D. D. & Olivier, C. Y. Aster leafhopper survival and reproduction, and Aster yellows transmission under static and fluctuating temperatures, using ddPCR for phytoplasma quantification. Sci. Rep. 8, 227 (2018).

22. Plante, N. et al. Leafhoppers as markers of the impact of climate change on agriculture. Cell Rep. Sustain. 1, 100029 (2024).

23. Baker, M. B., Venugopal, P. D. & Lamp, W. O. Climate change and phenology: *Empoasca fabae* (Hemiptera: Cicadellidae) migration and severity of impact. PLoS ONE 10, e0124915 (2015).

24. Taylor, R. A. J. & Reling, D. Preferred wind direction of long-distance leafhopper (*Empoasca fabae*) migrants and its relevance to the return migration of small insects. J. Anim. Ecol. 55, 1103–1114 (1986).

25. Taylor, R. A. J. & Shields, E. J. Revisiting potato leafhopper, *Empoasca fabae* (Harris), migration: implications in a world where invasive insects are all too common. Am. Entomol. 64, 44–51 (2018).

26. Chasen, E. M., Dietrich, C., Backus, E. A. & Cullen, E. M. Potato Leafhopper (Hemiptera: Cicadellidae) ecology and integrated pest management focused on alfalfa. J. Integr. Pest Manag. 5, A1–A8 (2014).

27. Bertaccini, A. et al. Revision of the ‘*Candidatus* Phytoplasma’ species description guidelines. Int. J. Syst. Evol. Microbiol. 72 (2022).

28. Maejima, K., Oshima, K. & Namba, S. Exploring the phytoplasmas, plant pathogenic bacteria. J. Gen. Plant. Pathol. 80, 210–221 (2014).

29. Weintraub, P. G., Trivellone, V. & Krüger, K. The biology and ecology of leafhopper transmission of phytoplasmas 27-52 (ed. Bertaccini, A., Weintraub, P. G., Rao, G. P. & Mori, N.) (Springer Singapore, 2019).

30. Brochu, A.-S., Dionne, A., Fall, M. L. & Pérez-López, E. A decade of hidden phytoplasmas unveiled through citizen science. Plant Dis. 107, 3389–3393 (2023).

31. Olivier, C. Y., Séguin-Swartz, G., Galka, B. & Olfert, O. O. Aster yellows in leafhoppers and field crops in Saskatchewan, Canada, 2001–2008. Am. J. Plant Sci. Biotechnol. 5, 88–94 (2011).

32. Nault, L. R. Maize Bushy Stunt and Corn Stunt: A Comparison of Disease Symptoms, Pathogen Host Ranges, and Vectors. Phytopathology 70, 659–662 (1980).

33. Lee, I.-M. et al. New subgroup 16SrIII-Y phytoplasmas associated with false-blossom diseased cranberry (*Vaccinium macrocarpon*) plants and with known and potential insect vectors in New Jersey. Eur. J. Plant. Pathol. 139, 399–406 (2014).

34. Almeida Santos, A., Jacques, J., Plante, N., Fournier, V. & Pérez-López, E. Leafhoppers as vectors of phytoplasma diseases in Canadian berry crops: a review in the face of climate change. Ann. Entomol. Soc. Am. 117, 14–20 (2024)

35. Botti, S. & Bertaccini, A. Grapevine yellows in Northern Italy: molecular identification of Flavescence dorée phytoplasma strains and of Bois Noir phytoplasmas: Phytoplasmas in grapevine yellows. J. Appl. Microbiol. 103, 2325–2330 (2007).

36. Gajardo, A. et al. Phytoplasmas associated with grapevine yellows disease in Chile. Plant Dis. 93, 789–796 (2009).

37. Romero, B., Olivier, C., Wist, T. & Prager, S. M. Oviposition behavior and development of aster leafhoppers (Hemiptera: Cicadellidae) on Selected host plants from the Canadian Prairies. J. Econ. Entomol. 113, 2695–2704 (2020).

38. Olivier, C. Y., Lowery, D. T. & Stobbs, L. W. Phytoplasma diseases and their relationships with insect and plant hosts in Canadian horticultural and field crops. Can. Entomol. 141, 425–462 (2009).

39. Lenzi, P., Stoepler, T. M., McHenry, D. J., Davis, R. E. & Wolf, T. K. *Jikradia olitoria* (Hemiptera:Cicadellidae) transmits the sequevar NAGYIIIβ phytoplasma strain associated with North American grapevine yellows in artificial feeding assays. J. Insect Sci. 19, 1 (2019).

40. Arocha-Rosete, Y. et al. Surveys reveal a complex association of phytoplasmas and viruses with the blueberry stunt disease on Canadian blueberry farms. Ann. Appl. Biol. 174, 142–152 (2019).

41. Hammond, C. et al. Detection of blueberry stunt phytoplasma in Eastern Canada using cpn60-based molecular diagnostic assays. Sci. Rep. 11, 22118 (2021).

42. Pérez-López, E., Omar, A. F., Al-Jamhan, K. M. & Dumonceaux, T. J. Molecular identification and characterization of the new 16SrIX-J and cpn60 UT IX-J phytoplasma subgroup associated with chicory bushy stunt disease in Saudi Arabia. Int. J. Syst. Evol. Microbiol. 68, 518–522 (2018).

43. Olivier, C. et al. Occurrence of phytoplasmas in leafhoppers and cultivated grapevines in Canada. Agric. Ecosyst. Environ. 195, 91–97 (2014).

44. Pagé, D. Acquisition de connaissances sur les phytoplasmes dans la culture du bleuetier en corymbe. https://www.agrireseau.net/petitsfruits/documents/Rapport_IRDA_2013-06-03_phytoplasme_bleuetier-final.pdf (2013).

45. Chiykowski, L. N. *Endria inimica* (Say), a new leafhopper vector of a celery-infecting strain of aster-yellows virus in barley and wheat. Can. J. Bot. 41, 669–672 (1963).

46. Frazier, N. W. & Severin, H. H. P. Weed-host range of California aster yellows. Hilgardia 16, 619–650 (1945).

47. Gilmer, R. M. Insect transmission of X-disease virus in New York. Plant Dis. Rep. 38, 628–629 (1954).

48. Esau, K., Magyarosy, A. C. & Breazeale, V. Studies of the mycoplasma-like organism (MLO) in spinach leaves affected by the aster yellows disease. Protoplasma 90, 189–203 (1976).

49. Gilmer, R. M., Palmiter, D. H., Schaefers, G. A. & McEwen, F. L. Leafhopper transmission of X-disease virus of stone fruits in New York. (New York, 1966).

50. Mori, N. et al. Experimental transmission by *Scaphoideus titanus* Ball of two Flavescence doree-type phytoplasmas. VITIS - J. Grapevine Res. 41, 99–102 (2002).

51. Phillips, S. J., Dudík, M. & Schapire, R. E. Maxent software for modeling species niches and distributions (Version 3.4.1). http://biodiversityinformatics.amnh.org/open_source/maxent/ (2017).

52. Kass, J. M., et al. *wallace* 2: a shiny app for modeling species niches and distributions redesigned to facilitate expansion via module contributions. Ecography 2023, e06547 (2023).

53. Brown, J. L. SDMtoolbox: a python-based GIS toolkit for landscape genetic, biogeographic and species distribution model analyses. Methods Ecol. Evol. 5, 694–700 (2014).

54. Elith, J. et al. Novel methods improve prediction of species’ distributions from occurrence data. Ecography 29, 129–151 (2006).

55. Phillips, S. J., Anderson, R. P., Dudík, M., Schapire, R. E. & Blair, M. E. Opening the black box: an open-source release of Maxent. Ecography 40, 887–893 (2017).

56. Ross, H. H. A survey of the *Empoasca fabae* Complex (Hemiptera, Cicadellidae). Ann. Entomol. Soc. Am. 52, 304–316 (1959).

57. Bonfils, J. & Schvester, D. The leafhoppers (Homoptera: Auchenorrhyncha) and their relationship with vineyards in south-western France. Annales Epiphyties 11, 325–336 (1960).

58. Bertin, S. et al. Diffusion of the Nearctic leafhopper Scaphoideus titanus Ball in Europe: a consequence of human trading activity. Genetica 131, 275–285 (2007).

59. Saguez, J. et al. Diversity and abundance of leafhoppers in Canadian vineyards. J. Insect Sci. 14, 73 (2014).

60. Franklin, J. Mapping species distributions: spatial inference and prediction (ed. Franklin, J.) (Cambrige University Press, 2010).

61. Jarnevich, C. S., Stohlgren, T. J., Kumar, S., Morisette, J. T. & Holcombe, T. R. Caveats for correlative species distribution modeling. Ecol. Inform. 29, 6–15 (2015).

62. Fick, S. E. & Hijmans, R. J. WorldClim 2: new 1-km spatial resolution climate surfaces for global land areas. Int. J. Climatol. 37, 4302–4315 (2017).

63. Dormann, C. F. et al. Collinearity: a review of methods to deal with it and a simulation study evaluating their performance. Ecography 36, 27–46 (2013).

64. Kelley, M. et al. GISS-E2.1: Configurations and Climatology. J. Adv. Model Earth Syst. 12, e2019MS002025 (2020).

65. Calvin, K. et al. Climate Change 2023: Synthesis Report. Contribution of Working Groups I, II and III to the Sixth Assessment Report of the Intergovernmental Panel on Climate Change (ed. Core Writing Team., Lee, H. & Romero, J.) (IPCC, Geneva, Switzerland, 2023).

66. Hijmans, R. J. raster: Geographic Data Analysis and Modeling. R package version 3.6-23. https://CRAN.R-project.org/package=raster (2023).

67. Kass, J. M. et al. ENMeval 2.0: Redesigned for customizable and reproducible modeling of species’ niches and distributions. Methods Ecol. Evol. 12, 1602–1608 (2021).

68. Elith, J. et al. A statistical explanation of MaxEnt for ecologists: Statistical explanation of MaxEnt. Divers. Distrib. 17, 43–57 (2011).

69. Lobo, J. M., Jiménez-Valverde, A. & Real, R. AUC: a misleading measure of the performance of predictive distribution models. Glob. Ecol. Biogeogr. 17, 145–151 (2008).

70. Warren, D. L. & Seifert, S. N. Ecological niche modeling in Maxent: the importance of model complexity and the performance of model selection criteria. Ecol. Appl. 21, 335–342 (2011).

71. Allouche, O., Tsoar, A. & Kadmon, R. Assessing the accuracy of species distribution models: prevalence, kappa and the true skill statistic (TSS): Assessing the accuracy of distribution models. J. Appl. Ecol. 43, 1223–1232 (2006).

72. Luke, D. A User’s Guide to Network Analysis in R (ed. Luke, D.) (Springer International Publishing, 2015).

73. Dray, S. & Dufour, A.-B. The ade4 Package: Implementing the Duality Diagram for Ecologists. J. Stat. Soft. 22, (2007).

74. Csárdi, G., et al. igraph for R: R interface of the igraph library for graph theory and network analysis. R package version 1.6.0, https://zenodo.org/records/8240644 (2023).

75. Di Cola, V. et al. ecospat: an R package to support spatial analyses and modeling of species niches and distributions. Ecography 40, 774–787 (2017).

76. Oman, P. W. The Nearctic leafhoppers (Homoptera: Cicadellidae): a generic classification and check list. Mem. Entomol. Soc. Wash. 3, 1–253 (1949).

77. Krishnankutty, S. M., Dietrich, C. H., Dai, W. & Siddappaji, M. H. Phylogeny and historical biogeography of leafhopper subfamily Iassinae (Hemiptera: Cicadellidae) with a revised tribal classification based on morphological and molecular data. Syst. Entomol. 41, 580–595 (2016).

78. Sidumo, A. J., Shields, E. J. & Lembo, A. Estimating the potato leafhopper *Empoasca fabae* (Homoptera: Cicadellidae) overwintering range and spring premigrant development by using geographic information system. J. Econ. Entomol. 98, 757–764 (2005).

79. Whitcomb, R. F., Kramer, J., Coan, M. E. & Hicks, A. L. Ecology and Evolution of leafhopper-grass host relationships in North America (ed. Harris, K. F.) 121–178 (Springer New York, 1987).

80. Lawton, D. et al. Pest population dynamics are related to a continental overwintering gradient. Proc. Natl. Acad. Sci. U.S.A. 119, e2203230119 (2022).

81. Rossi, J.-P. & Rasplus, J.-Y. Climate change and the potential distribution of the glassy-winged sharpshooter (*Homalodisca vitripennis*), an insect vector of *Xylella fastidiosa*. Sci. Total Environ. 860, 160375 (2023).

82. Van Nieuwenhove, G. A., Frías, E. A. & Virla, E. G. Effects of temperature on the development, performance and fitness of the corn leafhopper *Dalbulus maidis* (DeLong) (Hemiptera: Cicadellidae). Agr. Forest Entomol. 18, 1–10 (2016).

83. Maw, H. E. L., Foottit, R., Hamilton, K. G. A. & Scudder, G. G. E. Checklist of the Hemiptera of Canada and Alaska - Family Cicadellidae (ed. Maw, H. E. L., Foottit, R., Hamilton, K. G. A. & Scudder, G. G. E.) 52–80 (NRC Research Press, 2000).

84. Dorey, J. B. et al. A globally synthesised and flagged bee occurrence dataset and cleaning workflow. Sci Data 10, 747 (2023).

85. Crossley, M. S. et al. Warmer temperatures trigger insecticide-associated pest outbreaks. Pest Manag. Sci. 80, 1008–1015 (2024).

86. Salar, P., Charenton, C., Foissac, X. & Malembic-Maher, S. Multiplication kinetics of Flavescence dorée phytoplasma in broad bean. Effect of phytoplasma strain and temperature. Eur. J. Plant Pathol. 135, 371–381 (2013).

87. Maggi, F., Galetto, L., Marzachì, C. & Bosco, D. Temperature-dependent transmission of *Candidatus* phytoplasma asteris by the vector leafhopper *Macrosteles quadripunctulatus* Kirschbaum. Entomologia 2, 202 (2014).

88. Sabato, E. O., Landau, E. C., Barros, B. A. & Oliveira, C. M. Differential transmission of phytoplasma and spiroplasma to maize caused by variation in the environmental temperature in Brazil. Eur. J. Plant. Pathol. 157, 163–171 (2020).

86. Brochu, A.-S., Methot, A., Breton, A.-M., Lacroix, C., Légaré, J.-P. & Pérez López, E. First report of a ‘Candidatus Phytoplasma asteris’ strain affecting lingonberry (*Vaccinium vitis-idaea*) and causing lingonberry stunt phytoplasma disease. New Dis Rep, 45, e12058 (2022).

87. Oliveira, C.M. & Frizzas, M.R. Eight Decades of Dalbulus maidis (DeLong & Wolcott) (Hemiptera, Cicadellidae) in Brazil: What We Know and What We Need to Know. Neotrop Entomol 51, 1–17 (2022).

88. Moya-Raygoza, G. Biological Control of the Leafhopper Dalbulus maidis in Corn Throughout the Americas: Interaction Among Phytoplasma-Insect Vector-Parasitoids. In: Olivier, C., Dumonceaux, T., Pérez-López, E. (eds) Sustainable Management of Phytoplasma Diseases in Crops Grown in the Tropical Belt. Sustainability in Plant and Crop Protection, vol 12. Springer, Cham. (2019).

89. Dalaisón-Fuentes, L. I., Pascual, A., Gazza, E., Welchen, E., Rivera-Pomar, R. & Ines-Catalano, M. Development of efficient RNAi methods in the corn leafhopper *Dalbulus maidis*, a promising application for pest control. Pest Manag Sci 78: 3108–3116 (2022).

