## Supplementary material for "Potential impact of climate change on Nearctic leafhopper distribution and richness in North America": Table S1-S2 and Figure S1

**Supplementary Table S1.** Pearson pair-wise test between 19 bioclimatic variables at a spatial resolution of 2.5 minutes after cropped for North America. Bio 1: Annual Mean Temperature; Bio 2: Mean Diurnal Range (Mean of monthly (max temp - min temp)); Bio 3: Isothermality (BIO2/BIO7) (×100); Bio 4: Temperature Seasonality (standard deviation ×100); Bio 5: Max Temperature of Warmest Month; Bio 6: Min Temperature of Coldest Month; Bio 7: Temperature Annual Range (BIO5-BIO6); Bio 8: Mean Temperature of Wettest Quarter; Bio 9: Mean Temperature of Driest Quarter; Bio 10: Mean Temperature of Warmest Quarter; Bio 11: Mean Temperature of Coldest Quarter; Bio 12: Annual Precipitation; Bio 13: Precipitation of Wettest Month; Bio 14: Precipitation of Driest Month; Bio 15: Precipitation Seasonality (Coefficient of Variation); Bio 16: Precipitation of Wettest Quarter; Bio 17: Precipitation of Driest Quarter; Bio 18: Precipitation of Warmest Quarter; Bio 19: Precipitation of Coldest Quarter.

| Variable | Bio1 | Bio10 | Bio11 | Bio12 | Bio13 | Bio14 | Bio15 | Bio16 | Bio17 | Bio18 | Bio19 | Bio2 | Bio3 | Bio4 | Bio5 | Bio6 | Bio7 | Bio8 | Bio9 |
|---|---|---|---|---|---|---|---|---|---|---|---|---|---|---|---|---|---|---|---|
| Bio1 | 1 | 0.91 | 0.97 | 0.19 | 0.54 | 0.04 | 0.3 | 0.41 | 0.09 | 0.02 | 0.23 | 0.46 | 0.75 | -0.8 | 0.86 | 0.94 | -0.7 | 0.37 | 0.7 |
| Bio10 | 0.91 | 1 | 0.78 | 0.21 | 0.5 | 0 | 0.17 | 0.39 | 0.07 | 0.14 | 0.07 | 0.45 | 0.46 | -0.4 | 0.98 | 0.75 | -0.3 | 0.5 | 0.49 |
| Bio11 | 0.97 | 0.78 | 1 | 0.17 | 0.51 | 0.09 | 0.34 | 0.37 | 0.12 | -0.1 | 0.33 | 0.41 | 0.85 | -0.9 | 0.71 | 0.99 | -0.8 | 0.26 | 0.77 |
| Bio12 | 0.19 | 0.21 | 0.17 | 1 | 0.75 | 0.84 | -0.6 | 0.83 | 0.9 | 0.8 | 0.79 | 0.26 | 0.08 | -0.1 | 0.18 | 0.15 | -0.1 | -0.1 | 0.34 |
| Bio13 | 0.54 | 0.5 | 0.51 | 0.75 | 1 | 0.43 | -0.1 | 0.95 | 0.5 | 0.62 | 0.56 | 0.43 | 0.41 | -0.4 | 0.47 | 0.46 | -0.3 | -0 | 0.56 |
| Bio14 | 0.04 | 0 | 0.09 | 0.84 | 0.43 | 1 | -0.8 | 0.48 | 0.98 | 0.55 | 0.84 | 0.08 | 0.02 | -0.1 | -0 | 0.1 | -0.2 | -0.2 | 0.3 |
| Bio15 | 0.3 | 0.17 | 0.34 | -0.6 | -0.1 | -0.8 | 1 | -0.1 | -0.8 | -0.5 | -0.5 | 0.15 | 0.48 | -0.4 | 0.17 | 0.31 | -0.3 | 0.09 | 0.18 |
| Bio16 | 0.41 | 0.39 | 0.37 | 0.83 | 0.95 | 0.48 | -0.1 | 1 | 0.56 | 0.75 | 0.6 | 0.4 | 0.31 | -0.3 | 0.36 | 0.32 | -0.2 | -0 | 0.46 |
| Bio17 | 0.09 | 0.07 | 0.12 | 0.9 | 0.5 | 0.98 | -0.8 | 0.56 | 1 | 0.63 | 0.84 | 0.1 | 0.02 | -0.1 | 0.04 | 0.13 | -0.1 | -0.2 | 0.31 |
| Bio18 | 0.02 | 0.14 | -0.1 | 0.8 | 0.62 | 0.55 | -0.5 | 0.75 | 0.63 | 1 | 0.38 | 0.18 | -0.1 | 0.19 | 0.13 | -0.1 | 0.19 | 0.19 | -0.1 |
| Bio19 | 0.23 | 0.07 | 0.33 | 0.79 | 0.56 | 0.84 | -0.5 | 0.6 | 0.84 | 0.38 | 1 | 0.14 | 0.33 | -0.4 | 0 | 0.34 | -0.5 | -0.4 | 0.61 |
| Bio2 | 0.46 | 0.45 | 0.41 | 0.26 | 0.43 | 0.08 | 0.15 | 0.4 | 0.1 | 0.18 | 0.14 | 1 | 0.6 | -0.3 | 0.57 | 0.3 | -0 | 0.13 | 0.36 |
| Bio3 | 0.75 | 0.46 | 0.85 | 0.08 | 0.41 | 0.02 | 0.48 | 0.31 | 0.02 | -0.1 | 0.33 | 0.6 | 1 | -0.9 | 0.44 | 0.82 | -0.8 | 0.06 | 0.72 |
| Bio4 | -0.8 | -0.4 | -0.9 | -0.1 | -0.4 | -0.1 | -0.4 | -0.3 | -0.1 | 0.19 | -0.4 | -0.3 | -0.9 | 1 | -0.3 | -0.9 | 0.96 | -0 | -0.8 |
| Bio5 | 0.86 | 0.98 | 0.71 | 0.18 | 0.47 | -0 | 0.17 | 0.36 | 0.04 | 0.13 | 0 | 0.57 | 0.44 | -0.3 | 1 | 0.66 | -0.2 | 0.51 | 0.43 |
| Bio6 | 0.94 | 0.75 | 0.99 | 0.15 | 0.46 | 0.1 | 0.31 | 0.32 | 0.13 | -0.1 | 0.34 | 0.3 | 0.82 | -0.9 | 0.66 | 1 | -0.9 | 0.26 | 0.76 |
| Bio7 | -0.7 | -0.3 | -0.8 | -0.1 | -0.3 | -0.2 | -0.3 | -0.2 | -0.1 | 0.19 | -0.5 | -0 | -0.8 | 0.96 | -0.2 | -0.9 | 1 | 0 | -0.7 |
| Bio8 | 0.37 | 0.5 | 0.26 | -0.1 | -0 | -0.2 | 0.09 | -0 | -0.2 | 0.19 | -0.4 | 0.13 | 0.06 | -0 | 0.51 | 0.26 | 0 | 1 | -0.2 |
| Bio9 | 0.7 | 0.49 | 0.77 | 0.34 | 0.56 | 0.3 | 0.18 | 0.46 | 0.31 | -0.1 | 0.61 | 0.36 | 0.72 | -0.8 | 0.43 | 0.76 | -0.7 | -0.2 | 1 |

**Supplementary Table S2**. Environmental niche similarity test between 14 Nearctic leafhopper species based on five environmental variables and the final occurrence records (Figure 2). P-values < 0.05 indicate that the two species' environmental niches are more similar than expected by chance (one-tailed test). Climate scenario: current.

| Species | *A. inimicus* | *C. humilis* | *C. geminatus* | *E. rosae* | *E. fabae* | *E. comes* | *E. vitis* | *E. ziczac* | *E. exitiosus* | *G. fennahi* | *J. olitoria* | *M. quadrilineatus* | *P. irroratus* | *S. titanus* |
|---|---|---|---|---|---|---|---|---|---|---|---|---|---|---|
| *Amplicephalus inimicus* | NA | | | | | | | | | | | | | |
| *Ceratagallia humilis* | 0.030 | NA | | | | | | | | | | | | |
| *Colladonus geminatus* | 0.090 | 0.020 | NA | | | | | | | | | | | |
| *Edwardsiana rosae* | 0.060 | 0.320 | 0.427 | NA | | | | | | | | | | |
| *Empoasca fabae* | 0.020 | 0.128 | 0.277 | 0.001 | NA | | | | | | | | | |
| *Erythroneura comes* | 0.030 | 0.030 | 0.020 | 0.227 | 0.019 | NA | | | | | | | | |
| *Erythroneura vitis* | 0.001 | 0.140 | 0.158 | 0.079 | 0.001 | 0.059 | NA | | | | | | | |
| *Erythroneura ziczac* | 0.001 | 0.001 | 0.020 | 0.168 | 0.019 | 0.019 | 0.05 | NA | | | | | | |
| *Exitianus exitiosus* | 0.080 | 0.188 | 0.257 | 0.217 | 0.010 | 0.099 | 0.019 | 0.168 | NA | | | | | |
| *Graphocephala fennahi* | 0.090 | 0.455 | 0.405 | 0.001 | 0.039 | 0.148 | 0.158 | 0.158 | 0.178 | NA | | | | |
| *Jikradia olitoria* | 0.001 | 0.227 | 0.050 | 0.039 | 0.010 | 0.05 | 0.010 | 0.079 | 0.019 | 0.010 | NA | | | |
| *Macrosteles quadrilineatus* | 0.001 | 0.050 | 0.207 | 0.060 | 0.040 | 0.04 | 0.001 | 0.001 | 0.118 | 0.039 | 0.039 | NA | | |
| *Paraphlepsius irroratus* | 0.001 | 0.050 | 0.118 | 0.089 | 0.019 | 0.010 | 0.010 | 0.010 | 0.099 | 0.108 | 0.029 | 0.001 | NA | |
| *Scaphoideus titanus* | 0.001 | 0.020 | 0.030 | 0.069 | 0.010 | 0.019 | 0.010 | 0.029 | 0.138 | 0.010 | 0.059 | 0.001 | 0.010 | NA |

**Supplementary Table S2**. Environmental niche similarity test between 14 Nearctic leafhopper species based on five environmental variables and the final occurrence records (Figure 2). P-values < 0.05 indicate that the two species' environmental niches are more similar than expected by chance (one-tailed test). Climate scenario: SSP1-2.6

| Species | *A. inimicus* | *C. humilis* | *C. geminatus* | *E. rosae* | *E. fabae* | *E. comes* | *E. vitis* | *E. ziczac* | *E. exitiosus* | *G. fennahi* | *J. olitoria* | *M. quadrilineatus* | *P. irroratus* | *S. titanus* |
|---|---|---|---|---|---|---|---|---|---|---|---|---|---|---|
| *Amplicephalus inimicus* | NA | | | | | | | | | | | | | |
| *Ceratagallia humilis* | 0.0495 | NA | | | | | | | | | | | | |
| *Colladonus geminatus* | 0.0198 | 0.0099 | NA | | | | | | | | | | | |
| *Edwardsiana rosae* | 0.0693 | 0.3465 | 0.1980 | NA | | | | | | | | | | |
| *Empoasca fabae* | 0.0198 | 0.0693 | 0.2178 | 0.1181 | NA | | | | | | | | | |
| *Erythroneura comes* | 0.0099 | 0.0495 | 0.0693 | 0.1287 | 0.0693 | NA | | | | | | | | |
| *Erythroneura vitis* | 0.0099 | 0.3069 | 0.0495 | 0.1980 | 0.0297 | 0.0099 | NA | | | | | | | |
| *Erythroneura ziczac* | 0.0099 | 0.0099 | 0.0693 | 0.1287 | 0.0594 | 0.0099 | 0.0198 | NA | | | | | | |
| *Exitianus exitiosus* | 0.1584 | 0.1882 | 0.2871 | 0.4851 | 0.0198 | 0.1882 | 0.0297 | 0.1584 | NA | | | | | |
| *Graphocephala fennahi* | 0.0594 | 0.3267 | 0.3663 | 0.0099 | 0.0198 | 0.0990 | 0.1089 | 0.2079 | 0.1386 | NA | | | | |
| *Jikradia olitoria* | 0.0198 | 0.2673 | 0.0693 | 0.0693 | 0.0099 | 0.0297 | 0.0396 | 0.0594 | 0.0297 | 0.0099 | NA | | | |
| *Macrosteles quadrilineatus* | 0.0099 | 0.0396 | 0.0198 | 0.0693 | 0.0297 | 0.0099 | 0.0099 | 0.0099 | 0.0495 | 0.0990 | 0.0297 | NA | | |
| *Paraphlepsius irroratus* | 0.0099 | 0.0495 | 0.0198 | 0.0594 | 0.0099 | 0.0099 | 0.0099 | 0.0099 | 0.0198 | 0.0495 | 0.0198 | 0.0099 | NA | |
| *Scaphoideus titanus* | 0.0099 | 0.0198 | 0.0198 | 0.0099 | 0.0099 | 0.0099 | 0.0099 | 0.0099 | 0.1088 | 0.0693 | 0.0198 | 0.0099 | 0.0099 | NA |

**Supplementary Table S2**. Environmental niche similarity test between 14 Nearctic leafhopper species based on five environmental variables and the final occurrence records (Figure 2). P-values < 0.05 indicate that the two species' environmental niches are more similar than expected by chance (one-tailed test). Climate scenario: SSP2-4.5.

| Species | *A. inimicus* | *C. humilis* | *C. geminatus* | *E. rosae* | *E. fabae* | *E. comes* | *E. vitis* | *E. ziczac* | *E. exitiosus* | *G. fennahi* | *J. olitoria* | *M. quadrilineatus* | *P. irroratus* | *S. titanus* |
|---|---|---|---|---|---|---|---|---|---|---|---|---|---|---|
| *Amplicephalus inimicus* | NA | | | | | | | | | | | | | |
| *Ceratagallia humilis* | 0.0495 | NA | | | | | | | | | | | | |
| *Colladonus geminatus* | 0.0495 | 0.0396 | NA | | | | | | | | | | | |
| *Edwardsiana rosae* | 0.0495 | 0.3267 | 0.2772 | NA | | | | | | | | | | |
| *Empoasca fabae* | 0.0099 | 0.1188 | 0.0990 | 0.0990 | NA | | | | | | | | | |
| *Erythroneura comes* | 0.0099 | 0.0198 | 0.0396 | 0.1089 | 0.0297 | NA | | | | | | | | |
| *Erythroneura vitis* | 0.0099 | 0.4059 | 0.0891 | 0.1485 | 0.0099 | 0.0198 | NA | | | | | | | |
| *Erythroneura ziczac* | 0.0198 | 0.0099 | 0.0594 | 0.1089 | 0.0693 | 0.0099 | 0.0198 | NA | | | | | | |
| *Exitianus exitiosus* | 0.0119 | 0.2475 | 0.3267 | 0.4455 | 0.0198 | 0.1782 | 0.0297 | 0.2673 | NA | | | | | |
| *Graphocephala fennahi* | 0.1089 | 0.3465 | 0.3366 | 0.0099 | 0.0297 | 0.1782 | 0.1089 | 0.1089 | 0.1584 | NA | | | | |
| *Jikradia olitoria* | 0.0099 | 0.3168 | 0.0792 | 0.0591 | 0.0099 | 0.0495 | 0.0099 | 0.0297 | 0.0198 | 0.0198 | NA | | | |
| *Macrosteles quadrilineatus* | 0.0099 | 0.0495 | 0.0198 | 0.0198 | 0.0297 | 0.0198 | 0.0099 | 0.0099 | 0.0990 | 0.0693 | 0.0198 | NA | | |
| *Paraphlepsius irroratus* | 0.0099 | 0.1386 | 0.0396 | 0.0693 | 0.0297 | 0.0099 | 0.0099 | 0.0099 | 0.0594 | 0.0891 | 0.0099 | 0.0198 | NA | |
| *Scaphoideus titanus* | 0.0099 | 0.0396 | 0.0099 | 0.0099 | 0.0099 | 0.0099 | 0.010 | 0.0099 | 0.0693 | 0.0594 | 0.0099 | 0.0099 | 0.0099 | NA |

**Supplementary Table S2**. Environmental niche similarity test between 14 Nearctic leafhopper species based on five environmental variables and the final occurrence records (Figure 2). P-values < 0.05 indicate that the two species' environmental niches are more similar than expected by chance (one-tailed test). Climate scenario: SSP3-7.0.

| Species | *A. inimicus* | *C. humilis* | *C. geminatus* | *E. rosae* | *E. fabae* | *E. comes* | *E. vitis* | *E. ziczac* | *E. exitiosus* | *G. fennahi* | *J. olitoria* | *M. quadrilineatus* | *P. irroratus* | *S. titanus* |
|---|---|---|---|---|---|---|---|---|---|---|---|---|---|---|
| *Amplicephalus inimicus* | NA | | | | | | | | | | | | | |
| *Ceratagallia humilis* | 0.0594 | NA | | | | | | | | | | | | |
| *Colladonus geminatus* | 0.0396 | 0.0099 | NA | | | | | | | | | | | |
| *Edwardsiana rosae* | 0.0891 | 0.3069 | 0.3564 | NA | | | | | | | | | | |
| *Empoasca fabae* | 0.0297 | 0.1089 | 0.1980 | 0.0891 | NA | | | | | | | | | |
| *Erythroneura comes* | 0.0297 | 0.0198 | 0.0198 | 0.1881 | 0.0198 | NA | | | | | | | | |
| *Erythroneura vitis* | 0.0099 | 0.3564 | 0.0792 | 0.2475 | 0.0099 | 0.0099 | NA | | | | | | | |
| *Erythroneura ziczac* | 0.0099 | 0.0099 | 0.0198 | 0.1386 | 0.0495 | 0.0099 | 0.0099 | NA | | | | | | |
| *Exitianus exitiosus* | 0.0792 | 0.2673 | 0.2178 | 0.4752 | 0.0297 | 0.2079 | 0.0099 | 0.2574 | NA | | | | | |
| *Graphocephala fennahi* | 0.0297 | 0.3069 | 0.3663 | 0.0297 | 0.0495 | 0.1089 | 0.0396 | 0.0693 | 0.0792 | NA | | | | |
| *Jikradia olitoria* | 0.0099 | 0.2871 | 0.0891 | 0.0396 | 0.0099 | 0.0792 | 0.0198 | 0.0297 | 0.0297 | 0.0099 | NA | | | |
| *Macrosteles quadrilineatus* | 0.0099 | 0.0693 | 0.0297 | 0.0396 | 0.0297 | 0.0099 | 0.0099 | 0.0396 | 0.1287 | 0.0495 | 0.0198 | NA | | |
| *Paraphlepsius irroratus* | 0.0099 | 0.1188 | 0.0198 | 0.0693 | 0.0099 | 0.0099 | 0.0099 | 0.0099 | 0.0297 | 0.0297 | 0.0198 | 0.0099 | NA | |
| *Scaphoideus titanus* | 0.0099 | 0.0297 | 0.0198 | 0.0396 | 0.0198 | 0.0099 | 0.0099 | 0.0099 | 0.0495 | 0.0693 | 0.0099 | 0.0099 | 0.0099 | NA |

**Supplementary Table S2**. Environmental niche similarity test between 14 Nearctic leafhopper species based on five environmental variables and the final occurrence records (Figure 2). P-values < 0.05 indicate that the two species' environmental niches are more similar than expected by chance (one-tailed test). Climate scenario: SSP5-8.5.

| Species | *A. inimicus* | *C. humilis* | *C. geminatus* | *E. rosae* | *E. fabae* | *E. comes* | *E. vitis* | *E. ziczac* | *E. exitiosus* | *G. fennahi* | *J. olitoria* | *M. quadrilineatus* | *P. irroratus* | *S. titanus* |
|---|---|---|---|---|---|---|---|---|---|---|---|---|---|---|
| *Amplicephalus inimicus* | NA | | | | | | | | | | | | | |
| *Ceratagallia humilis* | 0.0495 | NA | | | | | | | | | | | | |
| *Colladonus geminatus* | 0.0396 | 0.0099 | NA | | | | | | | | | | | |
| *Edwardsiana rosae* | 0.1287 | 0.3363 | 0.2970 | NA | | | | | | | | | | |
| *Empoasca fabae* | 0.0099 | 0.0990 | 0.2079 | 0.0693 | NA | | | | | | | | | |
| *Erythroneura comes* | 0.0198 | 0.0099 | 0.0198 | 0.2079 | 0.0198 | NA | | | | | | | | |
| *Erythroneura vitis* | 0.0099 | 0.2673 | 0.0495 | 0.2080 | 0.0198 | 0.0198 | NA | | | | | | | |
| *Erythroneura ziczac* | 0.0198 | 0.0099 | 0.0396 | 0.1287 | 0.0594 | 0.0099 | 0.0198 | NA | | | | | | |
| *Exitianus exitiosus* | 0.0693 | 0.2376 | 0.2673 | 0.4752 | 0.0495 | 0.1485 | 0.0099 | 0.2376 | NA | | | | | |
| *Graphocephala fennahi* | 0.0990 | 0.3861 | 0.2673 | 0.0198 | 0.0297 | 0.1584 | 0.0792 | 0.1188 | 0.0990 | NA | | | | |
| *Jikradia olitoria* | 0.0198 | 0.3069 | 0.0891 | 0.0891 | 0.0198 | 0.0396 | 0.0297 | 0.0297 | 0.0693 | 0.0099 | NA | | | |
| *Macrosteles quadrilineatus* | 0.0099 | 0.0297 | 0.0396 | 0.0792 | 0.0396 | 0.0099 | 0.0099 | 0.0099 | 0.0693 | 0.0594 | 0.0099 | NA | | |
| *Paraphlepsius irroratus* | 0.0099 | 0.0594 | 0.0396 | 0.0693 | 0.0099 | 0.0198 | 0.0099 | 0.0099 | 0.0297 | 0.0495 | 0.0198 | 0.0099 | NA | |
| *Scaphoideus titanus* | 0.0099 | 0.0198 | 0.0099 | 0.0495 | 0.0099 | 0.0099 | 0.0099 | 0.0099 | 0.0495 | 0.0594 | 0.0198 | 0.0099 | 0.0099 | NA |

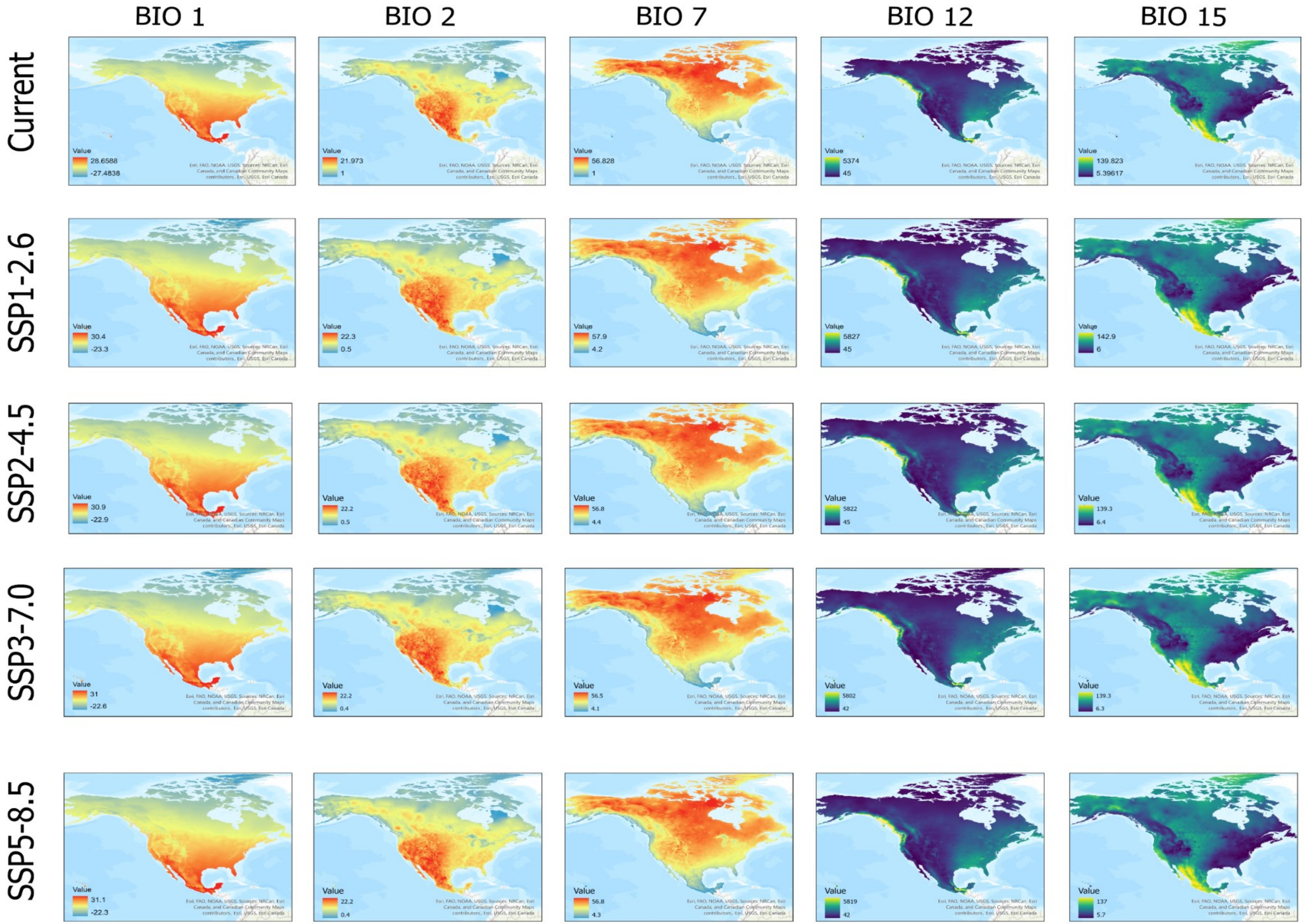


**Supplementary Figure 1**. Current (1970-2000) and future climate scenarios for 2041-2060 in North America. The SSP1-2.6 scenario limits global warming by 2100 to 2 °C, while the SSP2-4.5, SSP3-7.0, and SSP5-8.5 scenarios project values of 3 °C, 4 °C, and over 4 °C, respectively. BIO 1: Annual mean temperature; BIO 2: Mean diurnal range [mean of monthly(max temp – min temp)]; BIO 7: Temperature annual range; BIO 12: Annual precipitation; BIO 15:Precipitation seasonality (coefficient of variation).
